# Attachment of the RNA degradosome to the inner cytoplasmic membrane of *Escherichia coli* prevents wasteful degradation of rRNA intermediates in ribosome assembly

**DOI:** 10.1101/2022.06.14.496040

**Authors:** Lydia Hadjeras, Marie Bouvier, Isabelle Canal, Leonora Poljak, Quentin Morin-Ogier, Carine Froment, Odile Burlet-Schlitz, Lina Hamouche, Laurence Girbal, Muriel Cocaign-Bousquet, Agamemnon J. Carpousis

## Abstract

**Background:** RNase E has crucial roles in the initiation of mRNA degradation, the processing of ‘stable’ transcripts such as rRNA and tRNA, and the quality control of ribosomes. With over 20’000 potential cleavage sites, RNase E is a low specificity endoribonuclease with the capacity to cleave multiple times nearly every transcript in the cell. A large noncatalytic region in the C-terminal half of RNase E is the scaffold for assembly of the multienzyme RNA degradosome. The components of the RNA degradosome cooperate in the degradation of mRNA to oligoribonucleotides, which are then degraded to nucleotides by oligoribonuclease. Over the past decade, compelling evidence has emerged that the RNA degradosome is attached to the phospholipid bilayer of the inner cytoplasmic membrane by the Membrane Targeting Sequence (MTS), which is a 15-residue amphipathic alpha-helix located in the noncatalytic region of RNase E. Systematic proteomic analyses have identified RNase E as an inner membrane protein that can only be solubilized by disrupting the phospholipid bilayer with detergent. Important components of the mRNA degradation machinery are therefore membrane-attached. The reason for this cellular localization has until now been a mystery.

**Results:** We have constructed and characterized the *rneΔMTS* strain expressing ncRNase E (nucleo-cytoplasmic-RNase E), which is a soluble variant that is uniformly distributed in the interior of the cell. In the mutant strain, there is a slowdown in the rates of growth and mRNA degradation. Surprisingly, we have identified aberrant 20S and 40S ribosomal particles in the *rneΔMTS* strain that contain, respectively, precursors of 16S and 23S rRNA that have been ‘nicked’ by ncRNase E. Although intact ribosomes are resistant to RNase E cleavage *in vitro*, protein-free rRNA is readily degraded by RNase E. Partially unfolded ribosomes are susceptible to nicking by RNase E *in vitro*. We have mapped rRNA cleavage sites cRACE. *In vivo* and *in vitro* rRNA cleavages map to the same sites. The sequence of the cleavage sites matches the RNase E consensus sequence previously determined in a transcriptomic analysis that did not include rRNA. Construction of additional mutant strains demonstrated *in vivo* that fragments of 16S and 23S rRNA as well as a precursor of 5S rRNA are degraded in a pathway involving 3’ oligoadenylation and exonucleolytic digestion. A proteomic analysis showed that 17 small subunit proteins and 21 large subunit proteins are underrepresented in the 20S and 40S particles, respectively.

**Conclusions:** Ribosome biogenesis is a complex process involving co-transcriptional rRNA folding and r-protein binding in the nucleoid. Ribonucleoprotein intermediates are released from chromatin by RNase III cleavage. Maturation continues with the addition of ‘late’ proteins resulting in the compact rRNA structures found in mature 30S and 50S ribosomal subunits. Considering our experimental results, we propose that the physical separation of rRNA transcription in the nucleoid from the RNA degradosome on the inner cytoplasmic membrane protects intermediates in ribosome assembly from degradation. A corollary is that ribosome quality control normally occurs when defective ribosomal particles interact with the membrane-attached RNA degradosome. The rRNA degradation pathway described here is the same as described previously for RNase E-dependent degradation of mRNA. Since the pathway for rRNA degradation is the same as the pathway for mRNA degradation, the slowdown of mRNA degradation in the *rneΔMTS* strain could be due to competition by rRNA degradation. Since growth rate is limited by ribosome synthesis rate, the slow growth of the *rneΔMTS* strain is likely due to wasteful degradation of a proportion of newly synthesized rRNA. If r-proteins released by rRNA degradation are not recycled, then this would be an additional burden on cell growth. Avoiding a futile cycle in which rRNA intermediates in ribosome assembly are degraded likely explains why localization of RNase E homologues to the inner cytoplasmic membrane is conserved throughout the β- and γ-Proteobacteria.

**Importance:** In *E. coli*, transcription in the nucleoid, translation in the cytoplasm and initiation of mRNA degradation on the inner cytoplasmic membrane are physically separated. Despite the lack of internal membranes, this separation can be viewed as a compartmentalization of the bacterial cell. Our work shows that the inner membrane localization of the RNA degradosome restricts access of RNase E to intermediates in ribosome assembly. Thus, as in the eukaryotic cell, the architecture of the bacterial cell has an important role in the organization of cellular processes such as ribosome biogenesis, ribosome quality control, and mRNA degradation.

## Introduction

*E. coli* RNase E is the founding member of a large family of endoribonucleases that are widely distributed in bacteria and plants (Ait-Bara and Carpousis 2015; Ait-Bara et al. 2015). The N-terminal half of each subunit folds into a compact globular structure that forms the catalytic domain, while the C-terminal noncatalytic region is predominantly natively unstructured protein (Callaghan et al. 2004; Callaghan et al. 2005; Marcaida et al. 2006). The noncatalytic region has small motifs (15-40 residues) known as microdomains or SLiMs (<u>S</u>mall <u>Li</u>near <u>M</u>otifs), which serve as sites of interaction with proteins, RNA, and the phospholipid bilayer (Marcaida et al. 2006; Khemici et al. 2008; Ait-Bara and Carpousis 2015; Ait-Bara et al. 2015). The exoribonuclease PNPase, the glycolytic enzyme enolase, and the DEAD-box RNA helicase RhlB bind to RNase E microdomains to form the multienzyme RNA degradosome (Carpousis et al. 1994; Miczak et al. 1996; Py et al. 1996; Vanzo et al. 1998; Carpousis 2007). Another microdomain, known as the MTS (<u>M</u>embrane <u>T</u>argeting <u>S</u>equence), forms a 15-residue amphipathic alpha-helix that binds to phospholipid bilayers (Khemici et al. 2008; Strahl et al. 2015). Protein sequence comparisons have shown that RNase E homologs in the γ-Proteobacteria have a conserved N-terminal catalytic domain and a large natively unstructured C-terminal half with microdomains that include a conserved MTS (Ait-Bara et al. 2015). MTS-like microdomains have also been identified in RNase E homologs in β-Proteobacteria (Khemici et al. 2008).

Epifluorescence and super-resolution microscopy of live cells has shown that RNase E is localized to the periphery of the cell with no detectable protein in the nucleoid (Khemici et al. 2008; Strahl et al. 2015; Moffitt et al. 2016). Furthermore, systematic analyses of the inner membrane proteome have shown that RNase E is an inner membrane protein (IMP) that can only be solubilized by treatment with detergents that disrupt the phospholipid bilayer (Papanastasiou et al. 2013; Papanastasiou et al. 2016). For clarity, we will refer to wild type RNase E as imRNase E (inner-membrane-RNase E). As evidenced by epifluorescence and TIRF microscopy, imRNase E forms short-lived clusters (puncta) on the inner cytoplasmic membrane (Strahl et al. 2015; Hamouche et al. 2021a). RhlB and PNPase have been shown to display the same localization and dynamics as imRNase E thus confirming the association of these enzymes in live cells (Hamouche et al. 2021a). RNA degradosomes appear to move on the inner cytoplasmic membrane, but this movement could be an illusion due to the rapid formation and dissociation of puncta over a few seconds. Inhibition of transcription by rifampicin results in the depletion of mRNA, precursors of rRNA and tRNA, and the disassembly of RNA degradosome puncta suggesting that RNA substrate is required for clustering (Strahl et al. 2015; Hamouche et al. 2021a). However, recent work with kasugamycin, which inhibits the initiation of translation, also results in the disassembly of RNA degradosome puncta (Hamouche et al. 2021a). Although there is a low-level translation of leaderless mRNA in the presence of kasugamycin, velocity sedimentation analyses showed that polyribosomes are not formed (Kaberdina et al. 2009; Muller et al. 2016). Since transcription continues in the presence of kasugamycin, and ribosome-free mRNA and precursors of rRNA and tRNA continue to be synthesized, the formation of RNA degradosome puncta was therefore proposed to be due to an interaction with polyribosomes. Biochemical work has shown that the RNA degradosome binds ribosomes and polyribosomes thus supporting a direct interaction (Tsai et al. 2012). Taken together, the experimental work suggests that puncta are sites of mRNA degradation in which the initial step involves the capture of polyribosomes by the RNA degradosome.

Recent work suggests that the RNA degradosome can be displaced from the inner cytoplasmic membrane under conditions of stress. Upon transition from aerobic to anaerobic growth, cells filament and RNase E localizes to the interior of the cell in a diffuse pattern (Murashko and Lin-Chao 2017). Starvation of *E. coli* for a nitrogen source results in the formation of a single large focus of RNase E (McQuail et al. 2021). Treatment of cells with the protein synthesis inhibitor chloramphenicol results in the formation of foci of RNase E that are not attached to the inner cytoplasmic membrane (Hamouche et al. 2021a). These results suggest that stress-induced detachment of RNase E from the inner membrane could control RNase E accessibility to RNA substrates.

Although RNase E is an essential enzyme in *E. coli*, mutant strains encoding variants in which part or all of the C-terminal region is deleted are viable (Vanzo et al. 1998; Lopez et al. 1999; Ow et al. 2000; Leroy et al. 2002). Binding to the inner cytoplasmic membrane of *E. coli* is disrupted in mutant strains in which the amphipathic α-helix formed by the MTS has been mutated by amino acid substitution or deletion (Khemici et al. 2008). In the *rne(ΔMTS)* background, ncRNase E localizes uniformly to the interior of the cell (Khemici et al. 2008; Strahl et al. 2015; Moffitt et al. 2016). The *rne(ΔMTS)* strain exhibits a slow-growth phenotype, a slowdown of mRNA degradation, and accelerated degradation of ribosome-free mRNA (Hadjeras et al. 2019). Although a previous study proposed that membrane localization of RNase E preferentially destabilizes mRNA encoding inner membrane proteins (Moffitt et al. 2016), this preference was not considered statistically significant in a subsequent study (Hadjeras et al. 2019), which concluded that the slowdown in mRNA degradation is global. Here, we present experimental evidence suggesting that the slowdown in mRNA degradation is an indirect effect involving competition with the degradation of intermediates in ribosome assembly by ncRNase E.

Over the past two decades, evidence has emerged that ribosome assembly and rRNA processing are organized spatially (Bohne 2014). In *E. coli*, despite their separation by hundreds of kbp on the chromosome, most rRNA operons are in close proximity leading to the suggestion that there is a bacterial nucleolus (Gaal et al. 2016). Ribosomal RNA is transcribed as a single 30S precursor, which then undergoes extensive processing carried out by a battery of ribonucleases including RNase III and RNase E (Deutscher 2009; Shajani et al. 2011; Sulthana and Deutscher 2013; Jain 2020). EM imagining of chromosome spreads showed that ribosomal proteins bind to rRNA co-transcriptionally (Miller et al. 1970). Recent work has elucidated the structure of an rRNA transcription elongation complex that promotes co-transcriptional RNA folding and r-protein binding (Huang et al. 2020). Some maturation steps, such as processing by RNase III, take place on nascent rRNA in the nucleoid. RNA FISH (<u>F</u>luorescence In <u>S</u>itu <u>H</u>ybridization) showed that the 5’ leader sequence of 30S rRNA is localized to the nucleoid whereas the 17S precursor of 16S rRNA is localized in the cytoplasm, and that this separation is RNase III-dependent (Malagon 2013). These results show that early steps in ribosome biogenesis occur co-transcriptionally in the nucleoid.

Ribosome assembly is a complex multi-step process that requires the coordinated synthesis of rRNA and r-proteins (Shajani et al. 2011; Davis and Williamson 2017). *In vivo* kinetic analyses revealed the existence of two intermediates is 30S assembly (p_1_30S and p_2_30S) and three intermediates in 50S assembly (p_1_50S, p_2_50S and p_3_50S) (Lindahl 1973). The p_2_30S intermediate, which co-sediments with the mature 30S subunit, contains the full complement of 21 small subunit proteins and precursor 16S rRNA. The p_3_50S intermediate, which co-sediments with the mature 50S subunit, contains the full complement of 30 large subunit proteins and precursor 23S and 5S rRNA. The p_1_30S intermediate contains 10 small subunit proteins; p_1_50S, p_2_50S contain 16 and 24 large subunit proteins, respectively. In addition to r-protein binding, there are a large number of ribosome assembly factors including enzymes that modify rRNA, RNA helicases and protein chaperons. However, recent work employing quantitative mass spectrometry and single-particle cryoEM structural analyses has revealed that the ribosome assembly pathways are more complex than previously believed (Shajani et al. 2011; Davis and Williamson 2017).

Errors in ribosome biogenesis result in defective subunits that could interfere with translation. Recent work has shown that rRNA in defective ribosomal subunits is eliminated by degradation that is initiated by RNase E cleavage (Zundel et al. 2009; Basturea et al. 2011; Sulthana et al. 2016; Hamouche et al. 2021b). Here, we show that ncRNase E in the mutant *rneΔMTS* strain initiates rRNA degradation in assembly intermediates resulting in a wasteful cycle that slows cell growth and mRNA degradation. In the wild type strain, we propose that degradation of rRNA in defective ribosomal subunits is initiated by imRNase E cleavage at sites in 16S and 23S rRNA that are normally sequestered in intact, properly folded ribosomes.

## Results

### A genetic link between RNase E localization and ribosome biogenesis

We previously observed that the *rne*ΔMTS strain, which expresses ncRNase E, grows at approximately 80% of the rate of the isogenic *rne*^+^ strain expressing imRNase E (Hadjeras et al. 2019). To investigate the slow growth rate, we first analyzed cell shape and size. Visual inspection of the micrographs in Fig. 1A shows no obvious morphological difference between the *rne*ΔMTS and *rne*^+^ strains. This result suggests that the slower growth rate is not due to defective cell wall synthesis or cell division since the morphology is normal. Next, we measured cell size. In LB medium, there is a small decrease in cell length and width in the mutant strain that results in about a 10% decrease in cells size (Fig. 1B). Similar results were obtained in MOPS-glycerol medium although the difference in cell width is negligible. From these results, we conclude that the slower rate of growth of the *rne*ΔMTS strain correlates with a small decrease in cell size, which is consistent with known correlations between growth rate and cell size in *E. coli* (Zheng et al. 2020).

**Fig. 1.**
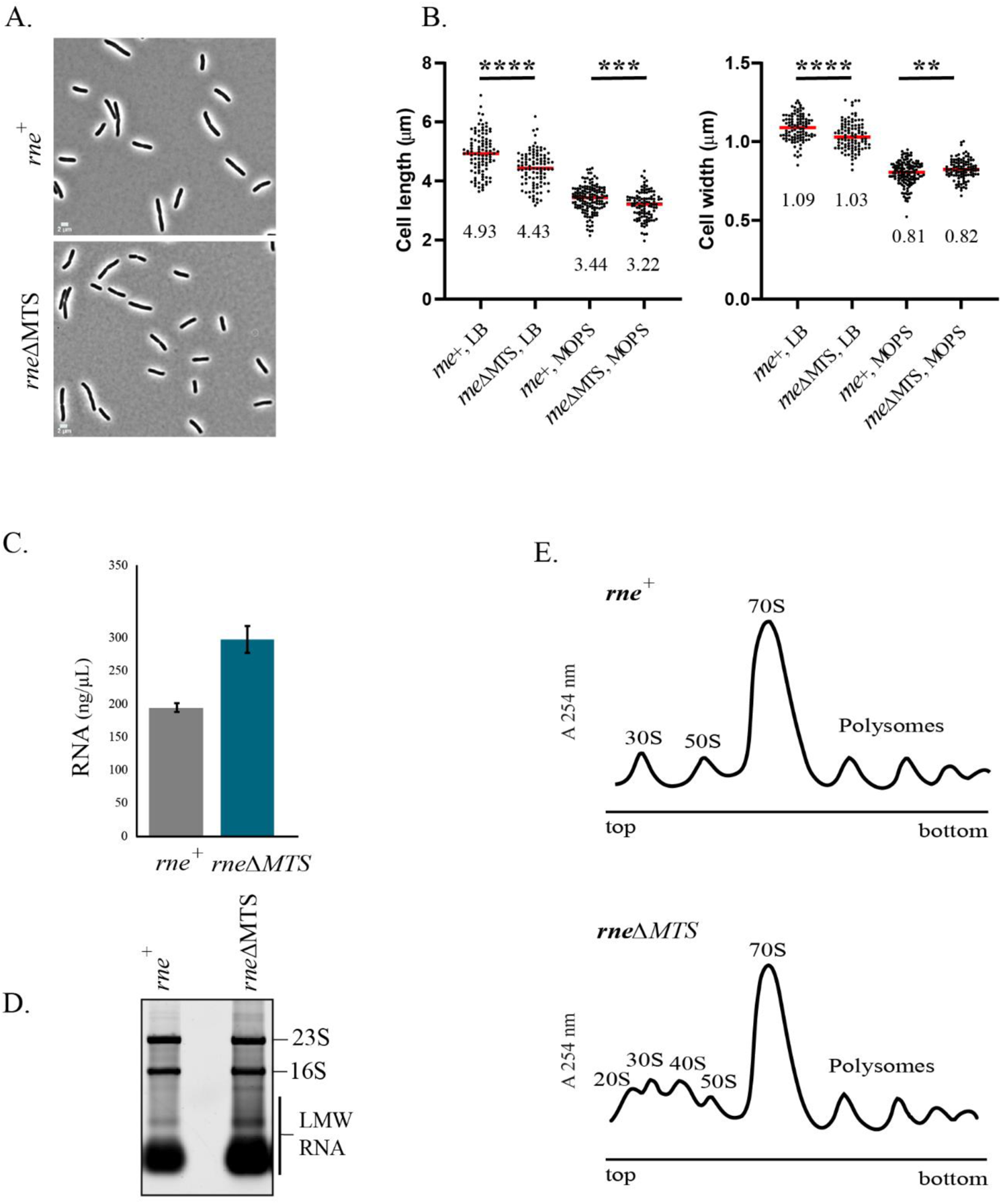
Cell size, RNA content and polysome profiles. A. Phase-contrast images. Micrographs of strains expressing either membrane-bound (*rne*^+^) or cytoplasmic (*rne*ΔMTS) RNase E were made at the same magnification.
B. Cell size. Lengths and widths were measured as described (Hamouche et al. 2021b). Scatter plots showing median cell length and width of *rne*^+^ and *rne*ΔMTS strains grown in either LB or MOPS media. Cells from two independent experiments (n>100) were analyzed by Image J using the MicrobeJ plugin. Median length and widths (µm) are shown below each plot. P-values were calculated using a parametric unpaired t-test (GraphPad Prism): **** = P < 0.0001; *** = 0.0001< P <0.001; ** = 0.001< P <0.01.
C. RNA yield. Cultures of the *rne*^+^ and *rne*ΔMTS strains were grown to OD_600_ =0.4 in LB medium. RNA was extracted from equal volumes of culture. Purified total RNA was eluted in equal volumes of water and concentrations were determined by UV absorption at 260 nm. Average and standard deviation of RNA concentration from three independent experiments are shown.
D. Ribosomal RNA levels. Equal volumes of total RNA (Fig. 1C) separated by agarose gel electrophoresis and staining with SybrGreen®. Levels of 16S and 23S are comparable in the two strains, whereas there is 30% more Low Molecular Weigh RNA (LMW RNA) in the *rne*ΔMTS strain as estimated by quantification of fluorescence levels (Image Lab, Biorad).
E. Polysome profiles. Clarified cell lysates prepared from equal volumes of cell cultures grown to OD_600_ =0.4 in LB medium were fractionated by velocity sedimentation on 10-40% sucrose gradients. Sedimentation is from left to right. Upper panel, *rne*^+^ strain; lower panel, *rne*ΔMTS strain. Peaks corresponding to 30S and 50S ribosomal subunits, 70S ribosomes and polysomes are indicated. 20S and 40S particles in the *rne*ΔMTS strain are indicated.

During the preparation of RNA for transcriptome analyses (Hadjeras et al. 2019), we noticed an increased level of Low Molecular Weight (LMW) RNA in the *rne*ΔMTS strain (Fig. S1). When we extracted total RNA from exponentially growing strains in LB, we consistently obtained about 50% more RNA from the *rne*ΔMTS strain (Fig. 1C). Since RNA was extracted from cultures grown to the same density (OD_600_=0.4) and there is only a small difference in cell size between the *rne*ΔMTS and *rne*^+^ strains, these results show a significant increase in total RNA levels in the *rne*ΔMTS mutant strain. We fractionated total RNA on an agarose gel by loading RNA extracted from equal volumes of cultures grown to the same density (Fig. 1D). The levels of 23S and 16S rRNA are comparable whereas the level of LMW RNA is about 30% higher in the mutant strain. These results show that the 50% increase in total RNA is at least partly due to an increase in LMW RNA. Although we have previously reported a slowdown in mRNA degradation in the *rne*ΔMTS strain (Hadjeras et al. 2019), it seems unlikely that the accumulation of mRNA degradation intermediates could by themselves explain the large increase in LMW RNA.

Comparable levels of 23S and 16S rRNA in the *rne*^+^ and *rne*ΔMTS strains strongly suggests that ribosome content in the mutant and wild type strains are comparable. Nevertheless, the slow growth phenotype could be due to a defect in translation resulting in lower protein synthesis rates. We therefore analyzed polyribosome profiles by velocity sedimentation on sucrose gradients to compare the level of 70S ribosomes to polyribosomes. Fig. 1E shows that the ratio of 70S ribosomes to polysomes is comparable between the two strains thus arguing against a defect in translation. However, the appearance of aberrant particles in the mutant strain with sedimentation coefficients of approximately 20S and 40S is striking. This result suggests a defect in ribosome assembly in the *rne*ΔMTS strain that could explain the slow growth phenotype.

### rRNA composition of 20S and 40S particles

To characterize the RNA composition of the 20S and 40S particles in the *rne*ΔMTS strain, sucrose gradient sedimentation was optimized to resolve the 20S to 70S region. RNA extracted from each sucrose gradient fraction was analyzed by slot blots probed with oligonucleotides specific to 17S, p16S, 16S, p23S, 23S and 5S rRNAs (Fig. 2A). For comparison, we have included an analysis of sucrose gradient fractions from the wild type strain. For both strains, as expected, 70S ribosomes contain mature rRNAs (fractions 28/29), whereas the 30S subunit (fractions 18/19/20) contains 17S, p16S and 16S rRNA and the 50S subunit (fractions 24/25/26) contains p23S, 23S and p5S rRNA. Analysis of polysome fractions by primer extension showed that they contain mature 5S, 16S and 23S rRNA (Fig. 2B). Detection of 17S, p16S, p23S and p5S rRNA in the wild type strain shows that a proportion of the 30S and 50S subunits, which correspond to the p_2_30S and the p_3_50S intermediates in ribosome assembly, are newly synthesized particles containing rRNA precursors (Lindahl 1973; Shajani et al. 2011). In the *rne*ΔMTS strain, the 20S particle contains the 17S and p16S precursors of 16S rRNA; the 40S particle contains the p23S and p5S precursors of 23S and 5S rRNA, respectively. The identification of rRNA precursors in the 20S and 40S particles was confirmed by primer extension (Fig. S2) and 5’ RACE (Fig. S3).

**Fig. 2.**
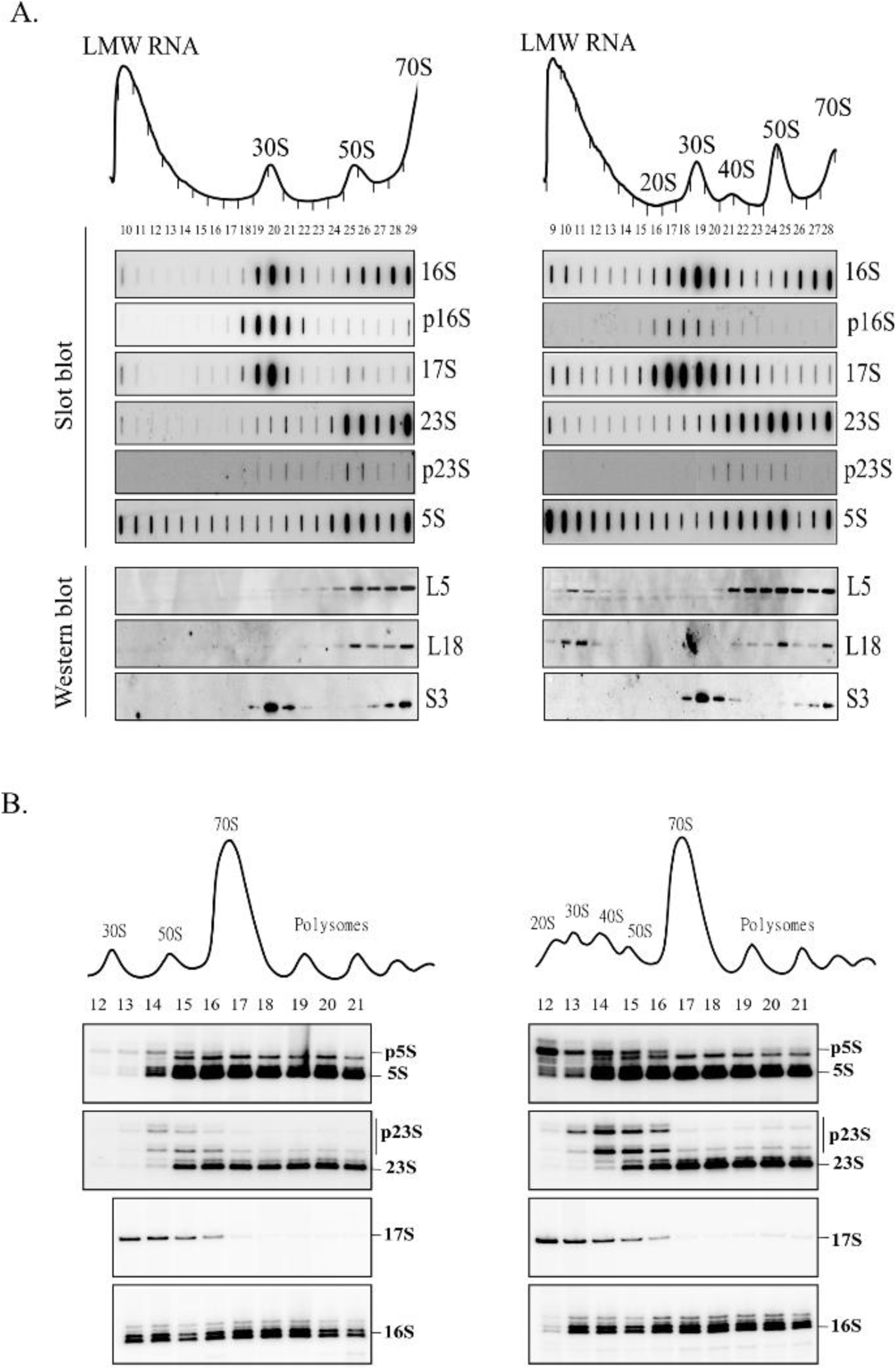
RNA content of ribosomal particles. Equal volume of clarified cell lysates from *rne*^+^ (left) and *rne*ΔMTS (right) strains were fractionated by velocity sedimentation. Sucrose gradient fraction numbers are indicated below the UV absorption profiles.

A. Conditions optimized for separation in the range of 20S to 70S. RNA from the sucrose gradient fractions was analyzed by slot blots using oligonucleotides specific to the RNA species indicated to the left of each panel. Protein from the sucrose gradient fractions was analyzed by Western blotting using antibodies against the ribosomal proteins indicated to the left of each panel.
B. Conditions optimized for separation of ribosomal subunits and polyribosomes. RNA from the sucrose gradient fractions was analyzed by primer extensions using [^32^P] end-labelled oligonucleotides specific to the 5’ ends of 5S, 23S, 17S and 16S rRNA. After extension by reverse transcriptase, the products were separated by denaturing gel electrophoresis. The 5’ end of mature rRNA and that of the prominent precursors are indicated to the right of each panel.

### Exonucleolytic degradation of rRNA by oligoadenylation and PNPase

The presence of p5S rRNA in the LMW (low molecular weight) region of the ribosome profile of the *rne*ΔMTS strain is striking (Fig. S2). In addition, p5S in LMW fractions co-sediments with L5 and L18 (Fig. 2A, Western blots), which are r-proteins known to bind to 5S rRNA (Korepanov et al. 2012). The co-sedimentation of p5S with L5/L18 in the LMW fractions is specific to the *rne*ΔMTS strain since they are almost undetectable in the *rne*^+^ strain. As a control, the sedimentation of S3, a 30S ribosomal protein, shows no differences in the *rne*^+^ and *rne*ΔMTS strains, suggesting low levels of most r-proteins in the LMW fractions. Taken together, these results suggest that a proportion of p5S rRNA that is complexed with the L5/L18 r-proteins fails to incorporate into mature 50S ribosomal subunit in the *rne*ΔMTS strain. Since p5S rRNA is the product of RNase E cleavage, these results also show that the defect in ribosome assembly is not due to a defect in RNase E processing of rRNA.

### Characterization of 5S* rRNA

Separation of total RNA on denaturing polyacrylamide gels, which resolve small RNA species in the range of 50 to 500 nt, revealed a prominent RNA in the *rne*ΔMTS strain, migrating slightly slower than 5S rRNA, that we named 5S* rRNA. (Fig. 3A, lanes 2). Primer extension of total RNA with an oligonucleotide specific to 5S rRNA detected the presence of mature 5S rRNA 5’ ends as well as species with 5’ end extension (Fig. 3B, lane 2). We have consistently seen two bands located between 5S and 5S* rRNA corresponding to species with 1 or 2 nt extensions, which agrees with work showing minor heterogeneity in the 5’ end of mature 5S rRNA (Feunteun et al. 1972; Jain 2020). We gel purified 5S rRNA from the *rne*^+^ and *rne*ΔMTS strains and 5S* rRNA from the *rne*ΔMTS strain and used RACE analyses to map the 5’ and 3’ ends of these molecules. A large proportion of the 5S rRNAs has a 5’ end corresponding to the mature molecule (Fig. S4A). In contrast, most of the 5S* rRNAs has a 5’ AUU extension that corresponds to the p5S precursor, which is generated by RNase E cleavage of 9S rRNA precursor. Analysis of 3’ ends showed that nearly all 5S rRNA molecules have a mature 3’ end whereas the 5S* molecules have heterogeneous 3’ ends (Fig. S4B). A large proportion of these molecules have the 3’ CAA extension that corresponds to the p5S rRNA precursor as well as untemplated oligo(A) additions ranging from 1 to 4 nt. The bands corresponding to 5S* rRNA (Fig. 3A, lanes 2) therefore corresponds to a mixture of p5S rRNA and oligoadenylated p5S rRNA. Oligo(A) additions are not detected in the *ΔpcnB* background, which lacks poly(A) polymerase activity (Fig. S4C). 3’ end analysis of RNA extracted from sucrose gradient fractions showed that p5S as well as p23S rRNA are oligoadenylated in the 40S particles from the *rne*ΔMTS strain whereas these RNAs were not oligoadenylated in 50S ribosomal subunits from the *rne*^+^ strain (Fig. S5).

**Fig. 3.**
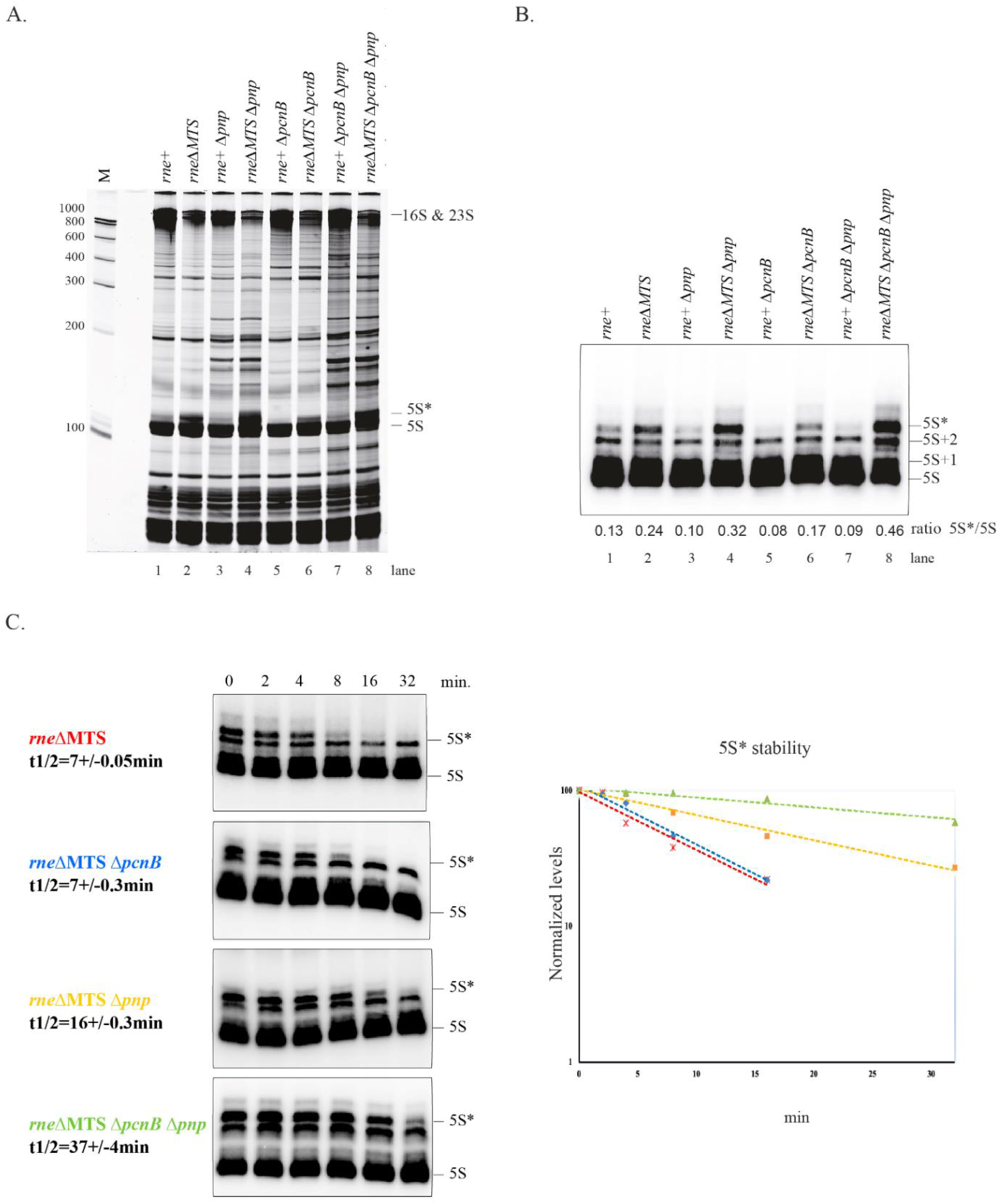
5S* rRNA is oligoadenylated form of p5S rRNA. A. Total RNA (10 µg) was extracted from strains that were grown in LB at 37 °C to OD_600_=0.4, separated on a denaturing polyacrylamide gel (10%, 7M urea) for 5h at 300V in 1x TBE, and then stained with SybrGreen dye.
B. Total RNA (1 µg) was analyzed by primer extension with a probe specific for 5S rRNA. The position of mature 5S rRNA and its precursors are indicated on the right. The levels of 5S and 5S* were quantified by phosphorimaging. The 5S*/5S ratio is indicated at the bottom of each lane.
C. The decay of 5S* rRNA was measured after the inhibition of transcription by rifampicin (left panel). Strains and half-lives are indicated to the left of each panel. Semi-log plot of quantification by phosphoimaging used to calculate half-lives (right panel). Mean half-live and standard deviation were determined from two or three independent experiments for each strain.

Since oligo(A) addition is indicative of 3’ exonucleolytic degradation, we compared the electrophoretic profiles of total RNA in the mutant strains lacking PNPase and poly(A) polymerase (Fig. 3A) and determined levels of 5S* rRNA by primer extension (Fig. 3B). Isogeneic *rne^+^* and *rne*ΔMTS strains were compared to reveal phenotypes specifically associated with the MTS deletion. In the *rne^+^* strain, there are trace amounts of 5S* rRNA. Levels expressed as the ratio of 5S*/5S shows that deletion of the genes encoding poly(A)polymerase and PNPase results in a large increase in 5S* rRNA. Deletion of the gene encoding PNPase alone also results in an increase in 5S* rRNA. Deletion of the gene encoding poly(A)polymerase results in the lower levels of 5S* rRNA, which is consistent with blockage of oligoadenylation. A kinetic analysis of 5S* rRNA decay after rifampicin treatment shows that p5S rRNA in the *rne*ΔMTS strain is degraded in a 3’ exonucleolytic pathway involving the activities of poly(A) polymerase and PNPase (Fig. 3C). Since the effect of deleting both enzymes is cumulative, the poly(A)polymerase-dependent pathway likely involves RNase R activity (see (Khemici and Carpousis 2004; Cheng and Deutscher 2005)).

## 16S and 23S rRNA are fragmented by ncRNase E

Since previous work has shown that RNase E has an essential role in ribosome quality control (Sulthana et al. 2016), we asked if 16S and 23S rRNA is fragmented in the 20S and 40S particles, respectively. To identify internal RNase E cleavages in 16S and 23S rRNA, we used an *exo^-^* strain background to knock down 3’ exonuclease activity and thereby increase the level of rRNA fragments. Although there are a large number of 3’ exonucleases in *E. coli*, RNase R and PNPase have a major role in the degradation of rRNA. Since inactivation of both genes encoding these enzymes is lethal, the *exo^-^* background combines a knockout of the *rnr* gene with the *pnp-200* allele, which expresses partially a partially inactive variant of PNPase (Cheng and Deutscher 2003). Fig. 4A shows sedimentation profiles of ribosomes from strains in which the Δ*rnr* and *pnp-200* alleles were moved into the *rne*^+^ and *rne*ΔMTS strains. Total RNA was extracted from each fraction of the gradient and then separated by gel electrophoresis. As expected for both strains, full-length 23S and 16S rRNAs are found in 70S ribosomes and are present in 50S and 30S subunits, respectively. In the *rne*ΔMTS strain, the 20S particle, which is essentially devoid of intact 16S rRNA, contains shorter RNA species that are about 1000 and 500 nt in length. The 40S particle contains 23S rRNA as well as shorter RNA species that are about 1700 and 1000 nt in length. In addition, fragments of about 500 nt are conspicuous in the LMW RNA fractions of the *rne*ΔMTS strain.

**Fig. 4.**
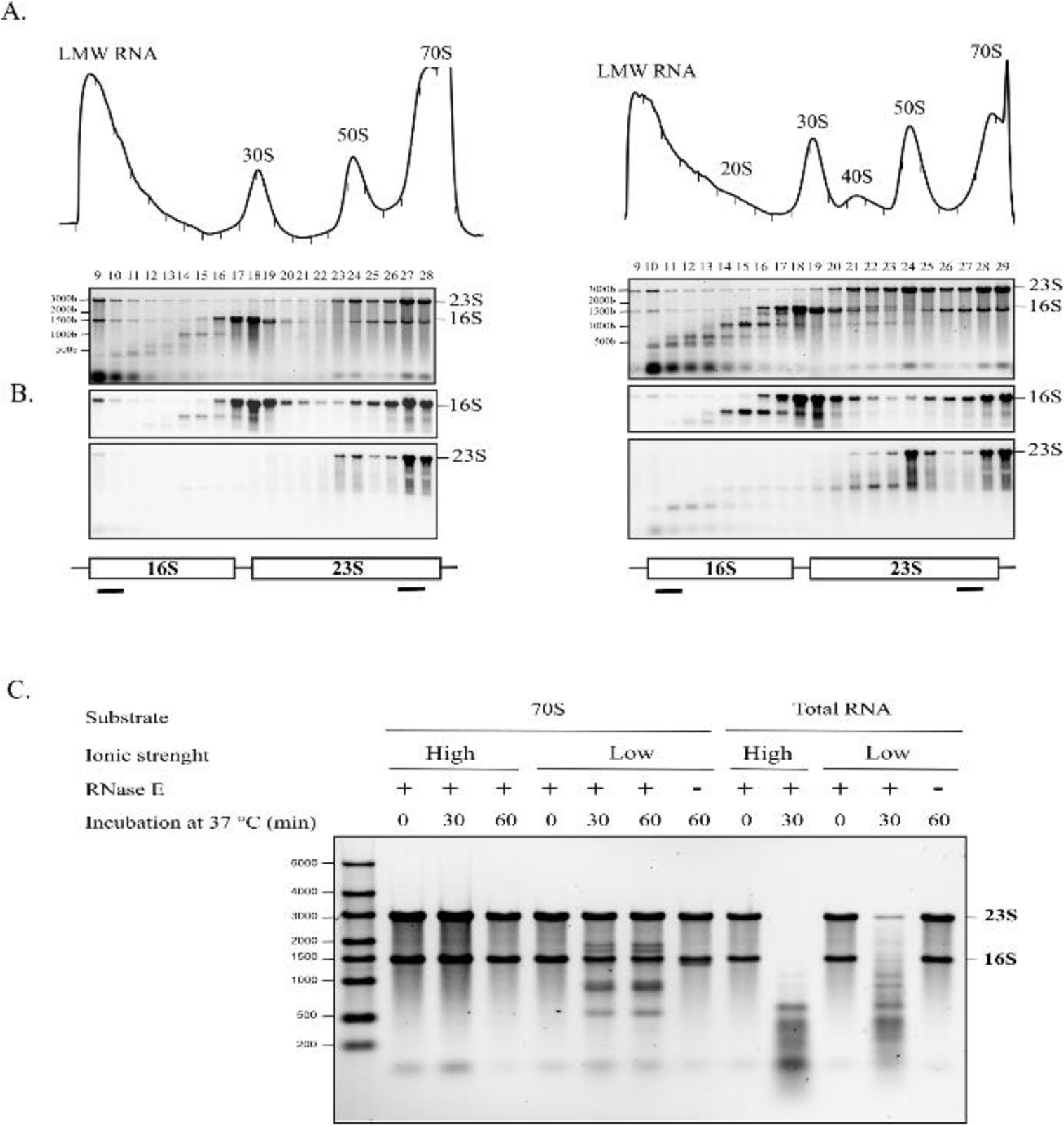
Identification of 16S and 23S rRNA fragments. A. Equal volumes of cell lysates from the *rne*^+^ *exo^-^* (left) and *rne*ΔMTS *exo^-^* (right) strains were separated by velocity sedimentation 5 to 20% sucrose gradients. RNA from each fraction was separated by gel electrophoresis.
B. Northern blots with probes specific to the 5’ end of 16S rRNA and the 3’ end of 23S rRNA as indicated in the diagram at the bottom of each panel.
C. Degradation of rRNA *in vitro*. RNase E cleavage assays were performed with purified 70S ribosomes or total RNA. A representative experiment is presented. After incubation at the indicated temperature and times, RNA was extracted, separated on 1% agarose gels and stained with SybrGreen. Each reaction contained 0.22 µM 70S ribosome or rRNA and 0.3 µM RNase E(1-598)-HIS6. Control lanes without RNase E (-) were also included. The position of the 23S and 16S rRNA are indicated to the right of the panel.

Northern blots of RNA from the sucrose gradients were probed with oligonucleotides specific to the 5’ end of 16S rRNA and the 3’ end of 23S rRNA (Fig. 4B). These blots show that the 1000 nt RNA fragment in the 20S particles contains the 5’ end of 16S rRNA and the 1700 nt RNA fragment in the 40S particle contains the 3’ end of 23S rRNA. These results strongly suggest that rRNA in the 20S and 40S particles is fragmented by endonucleolytic cleavage in the body of 16S and 23S rRNA.

### *In vitro* cleavage of ribosomal RNA by RNase E

We tested the activity of RNase E on ribosomes or rRNA *in vitro* (Fig. 4C). In a high ionic strength buffer, which is necessary for stability 70S ribosomes, rRNA is resistant to RNase E cleavage. Total RNA lanes at the right of the panel show that RNase E readily degrades protein-free rRNA in the high ionic strength buffer. Digestion, which results in a smear of fragments less than approximately 600 nt in length, shows that there rRNA has a large number of RNase E cleavage sites. Resistance of ribosomes to cleavage in the high ionic strength buffer shows that rRNA secondary and tertiary interactions and r-proteins protect rRNA from RNase E cleavage. The cleavage of protein-free rRNA by RNase E is slower in the low ionic strength buffer due to the limiting amount of Mg^++^, which is necessary for RNase E activity. In the low ionic strength buffer, rRNA in ribosomes is nicked by RNase E to give a series of fragments ranging from 500 to 2000 nt in length. From these results, we conclude that a subset of RNase E cleavage sites is accessible when the ribosome is partially unfolded in the low ionic strength buffer.

### Mapping RNase E cleavage sites in ribosomal RNA

Using cRACE (circular Rapid Amplification of cDNA Ends), we mapped 16S and 23S rRNA cleavage sites *in vivo* in the *rne*ΔMTS*-exo^-^* strain and *in vitro* using purified RNase E and ribosomes. The strategy employed in this analysis is described in Fig. S6. Table S1, which is a tabulation of the cRACE results, shows that the 3’ ends of in *vivo* fragments often contain noncoded oligo(A) additions. Fig. 5A and B are schematic diagrams indicating RNase E cleavage sites mapped by cRACE. The frequency (n) represents the number of times an end was sequenced. The color-coded key indicates cleavages that were mapped *in vivo*, *in vitro* or both *in vivo* and *in vitro*. Cleavages *in vivo* in the +22 to +32 of 16S rRNA results in a nested set of fragments with raggedy 3’ ends that are likely due to partial degradation by residual 3’ exonuclease activity in the *rne*ΔMTS*-exo^-^* strain. In Fig. 5A, many of the internal cleavages in 16S rRNA were detected both *in vivo* and *in vitro*, which validates the *in vitro* cleavage of partially unfolded ribosomes by RNase E as a faithful representation of cleavage *in vivo*. Fig. 5C shows the consensus sequence of rRNA cleavage sites that were mapped *in vivo*. The sequence is similar to the genome wide consensus obtained from 22,000 RNase E sites in *Salmonella* mRNA including the highly conserved U at position +2. (Chao et al. 2017b). These results are strong circumstantial evidence that ncRNase is responsible for cleavages of rRNA in the aberrant 20S and 40S particles.

**Fig. 5.**
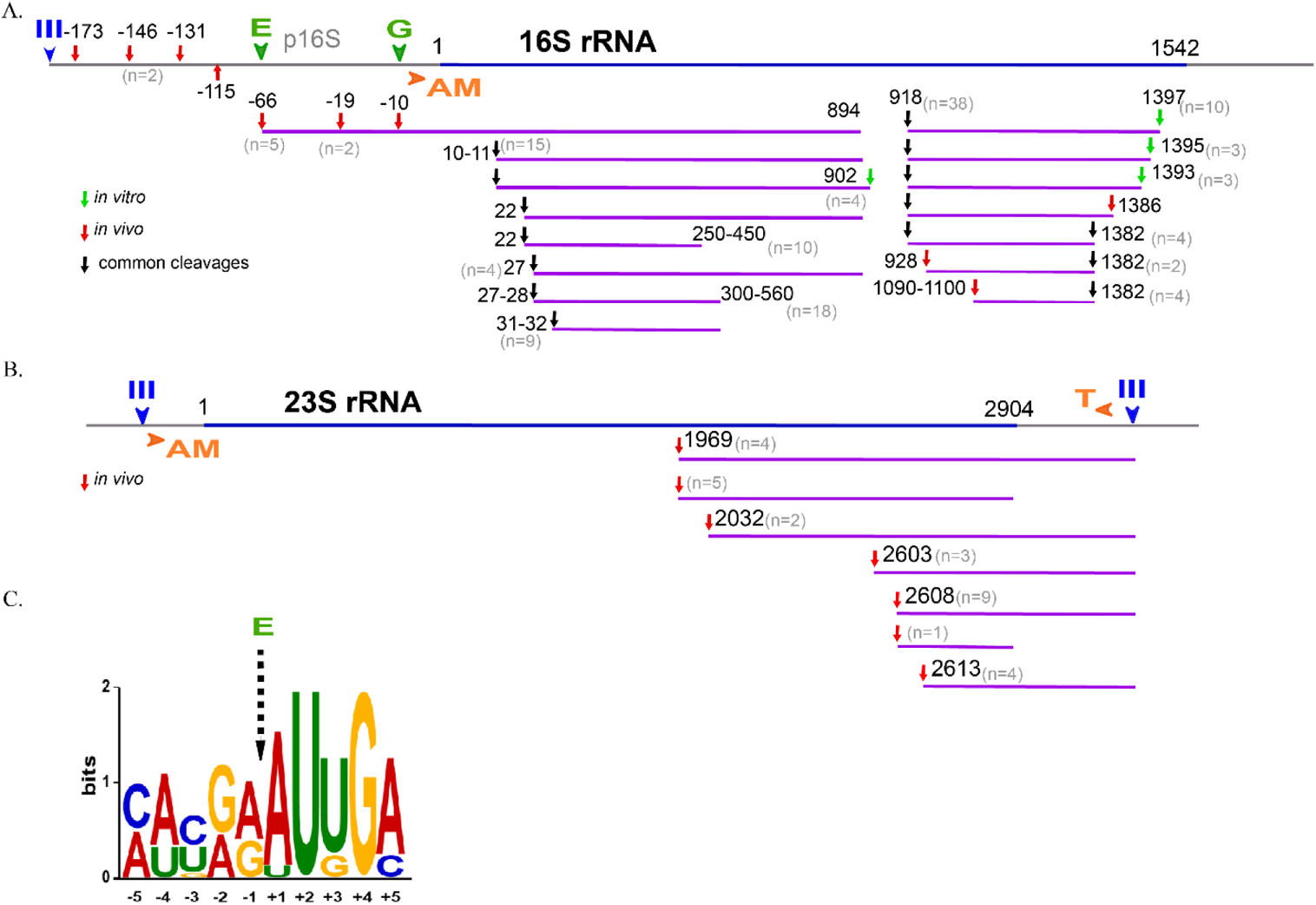
Mapping of RNase E cleavage sites. RNA fragments from the gels in Fig. 4 were extracted, purified, circularized and the region containing the 3’-5’ junction was PCR amplified (cRACE) as indicated in Fig. S6. After cloning the PCR fragments into a plasmid vector, the 3’-5’ junction was sequenced and the ends aligned with the sequence of 16S or 23S rRNA. In the diagrams representing 16S and 23S rRNA, III (blue), E (green), G (green), AM (orange) and T (orange) represent, respectively, rRNA processing sites for RNase III, RNase E and RNase G, which are endoribonucleases and RNase AM and RNase T, which are exoribonucleases that trim intermediates to the final mature species. **A. and B.** Identification of cleavage sites in 16S rRNA and 23S rRNA. The color-coded key indicates cleavages that were mapped *in vivo*, *in vitro* or both *in vivo* and *in vitro*. The number of times a site was sequenced is indicated (n). **C.** Consensus sequence of rRNA cleavage sites that were mapped *in vivo*.

### Proteomic analysis of the 20S and 40S ribosomal particles

We next analyzed protein content of the 20S and 40S particles from the *rne*ΔMTS strain. Proteins from sucrose fractions corresponding to these particles as well as to the 50S and 30S subunits of the *rne*^+^ and *rne*ΔMTS strains were extracted, digested with trypsin and then subjected to chromatography-tandem mass spectrometry (nanoLC-MS/MS), leading to the identification and quantification of 1286 proteins (detailed list in Table S2). To evaluate changes in protein compositions, pairwise comparisons based on MS intensity values were performed for each quantified protein, firstly, between *rne*^+^ and *rne*ΔMTS strains for 30S and 50S particles, secondly, between 20S and 30S particles as well as 40S and 50S particles in *rne*ΔMTS strain. Variant proteins were selected based on their significant protein abundance variations between the compared ribosomal particles (fold-change (FC) > 2 and < 0.5, and Student t test P < 0.05). Volcano plots in Fig. 6A and B show that composition of the 30S and 50S particles is globally the same in the two strains. The wild type 30S and 50S particles are enriched in integral and associated membrane proteins (pstG, secY, sdaC, bamD, murF, ubiG, proY, ccmE, gadC, mipA, ftsY and accY) (Karp et al. 2018; Keseler et al. 2021). Since the preparation of lysates for sucrose gradient analysis involves the use of sodium deoxycholate to solubilize membrane associated ribosomes, an interaction of imRNase E as part of detergent micelles containing other membrane proteins with the ribosomal subunits, could account for contamination with detergent solubilized membrane proteins. In *the rne*ΔMTS strain, 17 small subunit proteins and 21 large subunit proteins are significantly underrepresented in the 20S and 40S particles, respectively (Fig. 6C and D).

**Fig. 6.**
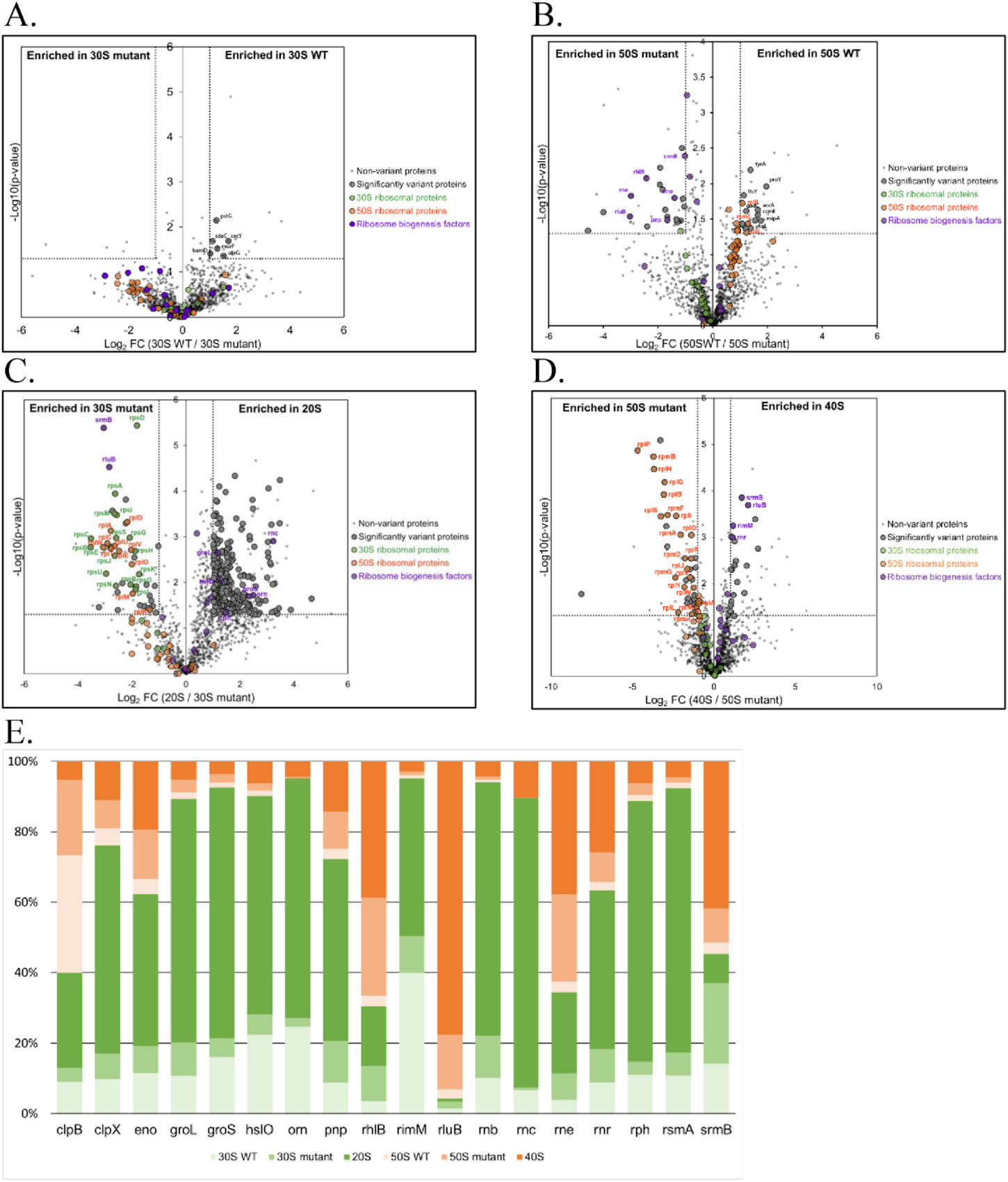
Protein composition of aberrant intermediates in ribosome assembly. Protein content of the ribosomal particles from *rne*^+^ and *rne*ΔMTS strains. Extracted proteins from sucrose gradient fractions were identified and quantified using a label-free quantitative mass spectrometry approach. Volcano plots showing significantly variant proteins (striped plots) in 30S particles from *rne*^+^ versus *rne*ΔMTS strains (**A.**), in 50S particles from *rne*^+^ versus *rne*ΔMTS strains (**B.**), in 20S versus 30S particles from *rne*ΔMTS strains (**C.**) and in 40S versus 50S particles from *rne*ΔMTS strains (**D.**), are presented. An unpaired bilateral student t-test with equal variance was used. Variant significance thresholds are represented by an absolute log2-transformed fold-change (FC) greater than 1 and a −log10-transformed (p-value) greater than 1.3 (see Materials and Methods). Small subunit proteins (green), large subunit proteins (orange) and ribosome biogenesis factors (purple) are indicated. **E**. Abundance levels of the quantified factors involved in ribosome biogenesis are represented as a percentage in 30S particle from *rne*^+^ strain (light green), in 30S particle from *rne*ΔMTS strain (medium green), in 20S particle (dark green), 50S particle from *rne*^+^ strain (light orange), in 50S particle from *rne*ΔMTS strain (medium orange) and in 40S particle (dark orange).

50S particles from the *rne*ΔMTS strain are enriched in ncRNase E, PNPase, RhlB and enolase, which are components of the RNA degradosome, as well as SrmB and RluB (Fig. 6B). SrmB is a DEAD-box RNA helicase that acts early in the assembly of the 50S subunit; RluB is a pseudouridine synthetase that acts late in the assembly of the 50S subunit (Karp et al. 2018; Keseler et al. 2021). Since it is likely that the gradient fractions analyzed here contain a mixture of particles (see Discussion), these results suggest that the 50S fraction from the *rne*ΔMTS contains a proportion of immature/defective particles whose degradation is initiated by the associated ncRNase E. 40S particles (Fig. 6D) are also enriched in SrmB and RluB as well as RNase R and RimM, which is a factor involved in the assembly of the 30S subunit (Karp et al. 2018; Keseler et al. 2021). The enrichment of RimM suggests a noncanonical interaction with aberrant 40S particles. The 20S particle is enriched for RNase III, RNase PH and oligoribonuclease (Fig. 6C, E) as well as the protein chaperones GroEL-GroES, which have been shown to have a role in assembly of the 50S ribosomal subunit (El Hage et al. 2001). Oddly, in the *rne*ΔMTS strain, the 30S fraction is enriched in a subset of large subunit proteins (Fig. 6C). A possible explanation for this result is that the 30S fraction in the *rne*ΔMTS strain is contaminated by slower sedimenting intermediates in the degradation of aberrant 40S particles. Proteins involved in ribosome assembly including enzymes that modify rRNA, ribonucleases, DEAD-box RNA helicases, the GroEL protein chaperone and the ClpXP protease are associated with the 20S and 40S particles, and most of these factors are underrepresented in the 30S and 50S ribosomal subunits (Fig. 6E). The underrepresented r-proteins and the associated ribosome assembly factors are supporting evidence for the conclusion that the 20S and 40S particles are aberrant dead-end intermediates in ribosome assembly.

## Discussion

Here we have shown that the *E. coli rneΔMTS* strain expressing ncRNase E has an abnormal ribosome profile with high levels of 20S and 40S particles. 5’ and 3’ end analysis showed that the particles contain precursors of 16S, 23S and 5S rRNA thus supporting the conclusion that they are intermediates in ribosome assembly as opposed to intermediates in the degradation of mature ribosomal particles. rRNA in the 20S and 40S particles is fragmented by ncRNase E cleavage within the 16S and 23S sequences. Mapping of ncRNase E cleavages in the 20S and 40S particles revealed sites whose sequences correspond to the consensus previously determined by genome-wide mapping of mRNA cleavages in *Salmonella* (Chao et al. 2017a). *In vitro* experiments with purified RNase E and ribosomes showed that properly folded ribosomes are resistant to RNase E cleavage whereas protein-free rRNA is readily degraded by RNase E. rRNA in ribosomes that are partially unfolded *in vitro* under low ionic strength conditions is cleaved by RNase E at sites that were mapped *in vivo*. From these results we conclude that rRNA cleavage sites in intact, properly folded ribosomes are sequestered by rRNA folding and r-protein binding.

In the *rneΔMTS* strain, fragments of 16S and 23 S rRNA as well as p5S rRNA have 3’ untemplated oligo(A) extensions. Oligoadenylated p5S rRNA migrates electrophoretically as a distinct species that we named 5S*. *In vivo* results with mutant strains showed that 3’ exonucleolytic degradation of 5S* rRNA involves the activities of PNPase and poly(A) polymerase. Measurements of 5S* degradation after rifampicin treatment showed an approximately 5-fold increase in half-life in a *pnp^-^*-*pcnB^-^* strain relative to the isogenic *rneΔMTS* control. It is also noteworthy that the exonucleolytic degradation pathway for 5S* rRNA is the same as previously described for several sRNA molecules as well as mRNA degradation intermediates containing REP elements (Khemici and Carpousis 2004; Cheng and Deutscher 2005; Andrade et al. 2012; Chen et al. 2021). Our velocity sedimentation results show that, in the Low Molecule Weight fraction at the top of the gradient, there are significant amounts of 5S rRNA precursors that co-sediment with proteins L5 and L18, which are known to bind to 5S rRNA (Korepanov et al. 2012). These results suggest that the p5S-L5-L18 complex accumulates as an intermediate in the *rneΔMTS* strain and that its failure to incorporate into the 50S ribosomal subunit triggers its degradation.

The 20S and 40S particles are, respectively, nominally equivalent to the p_1_30S intermediate, which sediments as a 21S particle, and the p_2_50S, which sediments as a 43S particle. Nevertheless, our proteomics analysis showing that 17 small subunit proteins and 21 large subunit proteins are underrepresented in the 20S and 40S particles, respectively, is inconsistent with their identification as *bone fide* intermediates in ribosome assembly. Furthermore, it is unlikely that the particles in the 20S and 40S sucrose gradient fractions are homogeneous in composition. Recent analysis of sucrose gradient fractions in the trailing edge of the 30S peak in a wild type strain showed that they contain a heterogeneous mixture of intermediates in ribosome assembly (Sashital et al. 2014). Similar results with intermediates in assembly of the 50S particle has led to the conclusion that ribosome assembly involves cooperative rRNA folding blocks that correspond to structural domains in the mature 30S and 50S ribosomal subunits, and that there are multiple parallel pathways leading to mature 30S and 50S ribosomal subunits (Davis et al. 2016; Davis and Williamson 2017).

Considering the large number of ncRNase E cleavages of rRNA that we have mapped in the *rneΔMTS* strain, we suspect that there are multiple pathways for the interference of ncRNase E with ribosome assembly. Although RNase E cleavage sites are single-stranded, the enzyme has the capacity to bind to structured RNA (Tsai et al. 2012; Bandyra et al. 2018). We therefore propose that ncRNase E competes directly with co-transcriptional r-protein binding resulting in misfolded intermediates lacking r-proteins. These dead-end intermediates are then cleaved by ncRNase E, which initiates their degradation. Although co-transcriptional interference with r-protein binding might be expected to trigger rho-dependent transcription termination, the rRNA transcription elongation complex is insensitive to rho-mediated termination (Condon et al. 1995; Huang et al. 2020). We believe that ncRNase E interference and rRNA cleavage are stochastic processes leading to a large number of different dead-end intermediates. The association of ribosome assembly factors with the 20S and 40S particles suggest that these factors are trying to ‘rescue’ dead-end intermediates. However, the degradation of rRNA in these particles suggests that the damage is mostly irreversible.

Our results strongly suggest that quality control of ribosomes is mediated by imRNase E. Fig. 7 is a cartoon depicting how the compartmentalization of the RNA degradosome to the inner cytoplasmic membrane protects partially unfolded intermediates in ribosome assembly from wasteful degradation. In this model, membrane attached RNA degradosomes are involved in the ‘trimming’ of 17S rRNA to p16S rRNA and 9S rRNA to p5S rRNA (Misra and Apirion 1979; Misra and Apirion 1980; Li et al. 1999). We propose that trimming of intermediates in ribosome assembly on the inner cytoplasmic membrane occurs after the subunits are properly folded and contain a full complement of r-proteins. This leads to the suggestion that the membrane attached RNA degradosome acts as a sensor that discriminates between properly folded, functional ribosomes and partially unfolded, inactive ribosomes that are degraded by the membrane-attached RNA degradosome. However, we believe that the interference of ncRNase E with ribosome assembly is likely to be mostly co-transcriptionally in the nucleoid and that normal ribosome quality control starts after intermediates are released from the nucleoid.

**Fig. 7.**
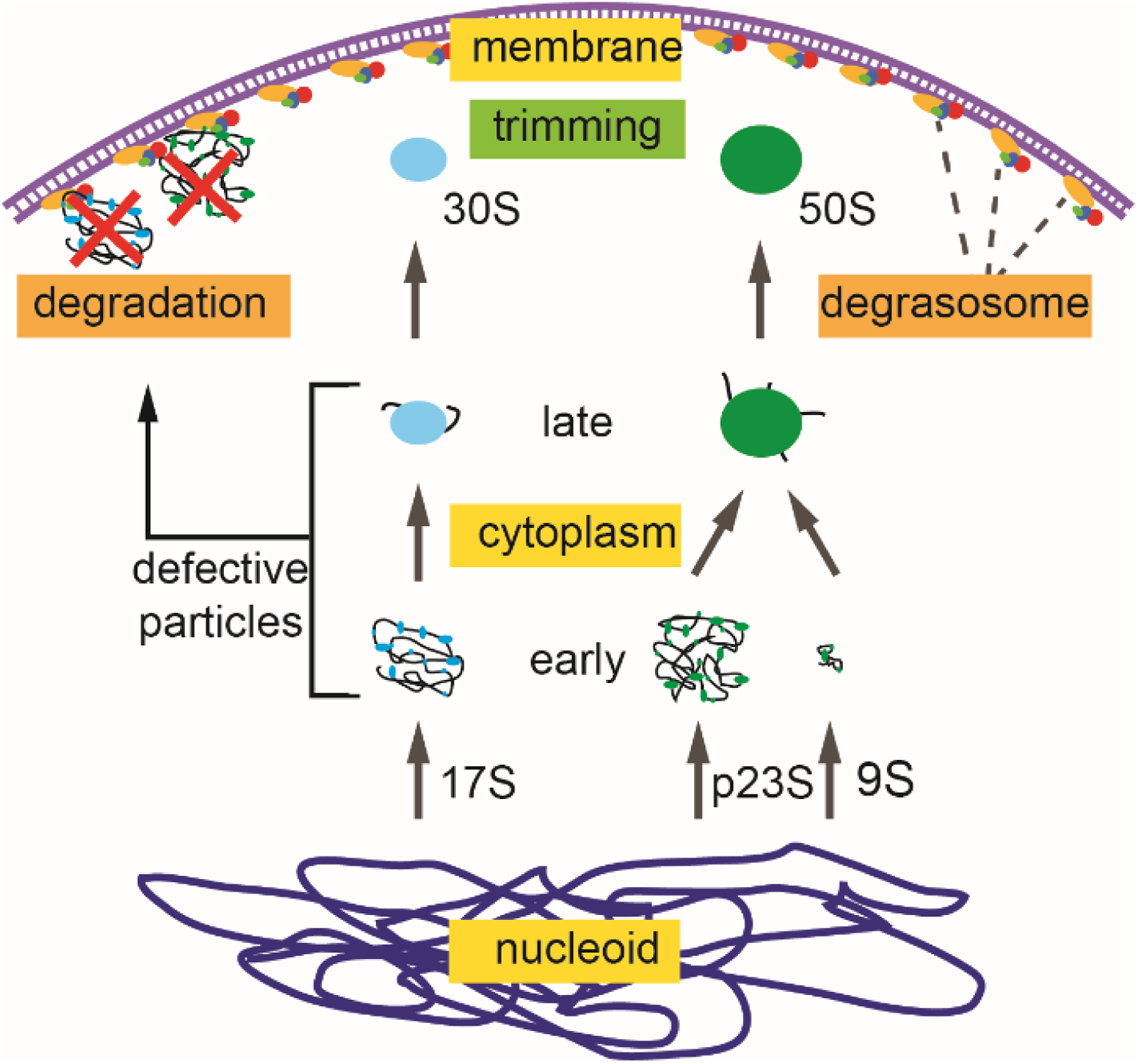
Quality control of ribosome assembly by the membrane attached RNA degradosome. Cartoon depicting the synthesis of rRNA in the nucleoid, the release of early intermediates in ribosome assembly from nucleoid, maturation of late intermediates in the cytoplasm, and trimming of 17S and 9S rRNA by the membrane attached RNA degradosome. Defective ribosomal particles are degraded by the membrane attached RNA degradosome. In this model, compartmentalization of ribosome assembly to the interior of the cell and the RNA degradosome on the inner cytoplasmic membrane shields intermediates in ribosome assembly from degradation thus avoiding wasteful turnover of rRNA. Defective particles can be either newly synthesized intermediates that have failed to properly fold or mature ribosomal subunits that are inactive (see Discussion).

Ribosome biogenesis is a major activity in growing cells. The time it takes for a cell to double is directly related to the time it takes to double ribosome content. Since rRNA synthesis is the limiting step in ribosome biogenesis (Paul et al. 2004), the wasteful degradation of rRNA likely explains the slower rate of growth of the *rneΔMTS* strain compared to the *rne^+^* strain (Khemici et al. 2008; Hadjeras et al. 2019). Enzymes involved in rRNA and mRNA degradation are the same (Carpousis et al. 2009; Zundel et al. 2009; Basturea et al. 2011; Sulthana et al. 2016; Hamouche et al. 2021b). Since recent work has shown the importance of competition between RNase E substrates in setting rates of mRNA degradation (Nouaille et al. 2017; Etienne et al. 2020), competition between rRNA and mRNA degradation could explain the global slowdown in mRNA degradation in the *rne(ΔMTS)* strain (Hadjeras et al. 2019). The work reported here shows that membrane attachment of RNase E as a component of the RNA degradosome is necessary to avoid a futile cycle of wasteful degradation of intermediates in ribosome assembly. Conservation of membrane-associated RNase E throughout the β- and γ-Proteobacteria is likely due to selective pressure to avoid interference with ribosome biogenesis.

RNase E homologues in the α-Proteobacteria lack identifiable MTS sequences and recent work, principally in *Caulobacter crescentus*, has shown that these enzymes are not attached to the inner cytoplasmic membrane. The RNA degradosome of *Caulobacter crescentus* is localized to the interior of the cell in condensates known as BR-bodies (<u>B</u>acterial <u>R</u>ibonucleoprotein-bodies) that have properties similar to eukaryotic stress granules and P-bodies (Al-Husini et al. 2018; Bayas et al. 2018). Assembly of BR-bodies is dynamic and requires RNA substrate as evidenced by rifampicin treatment. The endoribonuclease activity of RNase E is necessary for the disassembly of BR-bodies as evidenced by catalytically inactive variants of the enzyme. The intrinsically unstructured C-terminal region of RNase E, which is conserved in the α-Proteobacteria, is necessary and sufficient for BR-body formation. Characterization of the RNA content of *Caulobacter* BR-bodies showed that they are enriched in mRNAs and that rRNA and tRNA are excluded (Al-Husini et al. 2020). It was thus proposed that the *Caulobacter* BR-body is a compartment nucleated by the RNA degradosome in which mRNA is degraded. Selective permeability of the BR-body results in the enrichment of mRNA and mRNA decay intermediates thus increasing their concentration and driving degradation to nucleotides, which is important for maintaining nucleotide pools for transcription and DNA replication in growing cells. Importantly, BR-bodies form a compartment that is distinct from the nucleoid and cytoplasm. These results suggest that intermediates in ribosome assembly in *Caulobacter* are protected from nicking by *Caulobacter* RNase E due to the sequestration of the RNA degradosome into condensates that exclude ribosome precursors, ribosomes and polysomes.

Transcription and mRNA degradation in *Escherichia coli* and *Caulobacter crescentus* are physically separated in membraneless compartments. We therefore propose that these bacteria, which are separated by billions of years of evolution, use different strategies to achieve similar outcomes. Short-lived RNA degradosome puncta on the inner cytoplasmic membrane of *E. coli* are centers of mRNA degradation. The membrane attached RNA degradosome is also involved in the processing of rRNA and quality control of ribosomes. The physical separation of the RNA degradosome on the inner membrane from early steps in ribosome biogenesis in the nucleoid is necessary to prevent degradation of intermediates in ribosome assembly. The compartmentalization of RNA degradosomes in *Caulobacter* BR-bodies has functions similar to the membrane attachment of *E. coli* RNase E. BR-bodies are condensates in which the RNA degradosome and ribosome-free mRNA are concentrated thus driving degradation to nucleotides. BR-bodies exclude rRNA, ribosomes and polysomes thus segregating ribosome assembly from mRNA degradation. Compartmentalization of the mRNA degrading machinery in *E. coli* and *Caulobacter* is a fascinating example of evolution in which different cellular organizations result in solutions to similar problems involving the accessibility of RNA substrates to the RNA degradosome and the concerted degradation of mRNA to nucleotides.

## Material and Methods

### Bacterial strains and growth

The bacterial strains and oligonucleotides used in this study are included in the supplemental tables. Bacteria were grown either in LB or MOPS media prepared as described (Miller 1972; Neidhardt et al. 1974) at 180 rotations per minute (rpm) with normal aeration or agar plates at 37 °C. All mutant strains were constructed using the lambda-Red system as described in (Datsenko and Wanner 2000). After allele substitution into the chromosome using an antibiotic resistance cassette, the constructs were genetically purified by bacteriophage phage P1 transduction and the cassettes were removed using FLP recombinase resulting in an *frt* (FLP recognition target) scar All constructs were validated by sequencing PCR products amplified from chromosomal DNA.

### Cell dimension measurements

Samples for microscopy were prepared as in (Hamouche et al. 2021b). Briefly, bacterial strains were grown to OD_600_=0.5 at 37 °C with shaking in LB or MOPS medium supplemented with 0.5% glycerol and amino acids. Microscopy images were acquired on a Nikon Eclipse TI-E/B wide field epifluorescence microscope using phase contrast objective and were analyzed using Image J V.1.38 software. Statistical analysis and graphs were generated using GraphPad Prism, version 7.

### Polysome fractionation analysis

Polysomes fractionation analyses were performed as described (Charollais et al. 2003; Reyes-Lamothe et al. 2012; Qin and Fredrick 2013) with some modifications. Briefly, overnight cultures diluted in fresh LB medium were cultured at 37 ° C to an OD_600nm_ of 0.4. To stop bacterial growth and avoid ribosomes/polysomes dissociation, 40 OD_600_ equivalent units were harvested by fast-chilling by placing the cultures directly in a cold flask on an ice-water bath with shaking for 3 min. After centrifugation at 6000 g for 15 min at 4 °C (JA14 rotor-Beckman), the cell pellet is resuspended with cold lysis buffer (1mg/ml lysozyme, 10mM MgCl_2_, 60mM KCl, 10mM Tris-HCl pH8). For complete lysis, cells were subjected to two freeze-thaw cycles. After the second freeze-thaw cycle, 0.3% of sodium deoxycholate anionic detergent (D6750_SIGMA) was added to solubilize the membrane proteins and the lysate was clarified by centrifugation at 10000 g for 10min at 4 °C. To analyze polysome profiles, a constant volume of extract was layered onto an ultracentrifuge tube (tube 13.2mL-Beckman Coulter SW-41) containing a continuous 10-40% (w/v) sucrose gradient prepared in the following buffer: (10 mM MgCl_2_, 20mM Tris-HCl pH7.5, 100mM NH_4_Cl, 2mM dithiothreitol (DTT)) and centrifuged at 35000 rpm for 3h30 at 4 °C in an Optima XPN-80-Beckman Coulter ultracentrifuge. Sucrose gradients were analyzed on a density gradient fractionation system (ISCO UA-6 detector / Brandel Foxy Gradient) with continuous monitoring at 254nm, allowing the various ribosomal peaks to be resolved. To specifically analyze and resolve the ribosomal subunits, the extracts were layered onto a continuous 5-20% (w/v) sucrose gradient in the same buffer described above and centrifuged at 28600 rpm for 7h at 4 °C in a Beckman SW-41 rotor. The collected fractions were subjected to RNA and/or protein analyses.

### Total RNA extraction

2 to 4 OD_600_ of bacterial cell cultures were mixed with 0.2 volume of stop solution (ethanol: phenol 95:5 v/v) and snap-frozen in liquid nitrogen. Samples were thawed on ice, spun at 4000 rpm for 15 minutes at 4 °C and the cell pellet was dissolved in 1 mL TRIzol^R^ (Invitrogen, #15596026). An equal volume of ethanol was added to the mixture and total RNA was prepared using a Direct-zol™, RNA MiniPrep Plus kit (Zymo Research) following the manufacturer’s instructions and Dnase I digested using the Dnase provided in the same kit. RNA was eluted in 80 µl milliQ water (RNase-free) and RNA amount and purity were determined using a NanoDrop™ spectrophotometer.

### RNA isolation after fractionation on sucrose gradients

RNAs were extracted from sucrose gradient fractions by adding one volume of TRIzol^R^ and by using, the Direct-zol™ RNA Miniprep Plus kits (ZYMO RESEARCH, #R2072). The RNAs were eluted in 60µl of milliQ water (RNase-free) and subjected to DNase I digestion. After purification, the same amounts of RNA, unless indicated elsewhere, were used to perform primer extension analyses on the rRNAs as described below. Moreover, the same volume of RNA from sucrose fractions was also separated on native agarose gels (1%) (that do not contain formaldehyde), in 1X TBE buffer (10X TBE: 890mM Tris base, 890mM Boric acid, 20mM EDTA) for 3h30 at 50V. After electrophoresis, the gels were either subjected to Northern blotting as described below or stained with SYBR™ Safe stain (Invitrogen).

### Primer extension analysis

2 pmol of 5’end labelled primer (primers specific to 5S, 16S and 23S rRNA) and 0.25-1µg of RNA (total RNA or RNA extracted from sucrose gradient fractions) were denatured together in water for 5 min at 65 °C, and immediately quenched on ice for 5 min. 50 0µM dNTPs, 1x first strand buffer, 5mM DTT, 1 U/µl RNase inhibitor (ThermoScientific) and 1µl Superscript III reverse transcriptase (200U, Invitrogen) were added to the denatured RNA and primer (20µl reaction). Primer extension was allowed for 50 min at 55 °C. After heat inactivation of the reverse transcriptase for 5 min at 85 °C, samples were treated with 2U RNase H (Thermo Scientific) at 37 °C for 20 min. 2-5µl of the resulting reaction were mixed with an RNA loading dye and resolved on a 6% PAA, 7M urea sequencing gel along with the sequencing ladder. The sequencing ladder was obtained on a plasmid containing the 9S coding gene (*rrfB*) using USB® Sequenase ™ version 2.0 DNA polymerase (Affymetrix) following the supplier’s instructions. cDNA signals were visualized on a phosphoimager (Typhoon Trio-Amersham-Bioscience) and band intensities were quantified using MultiGauge software (Fujifilm).

### Northern blotting

5 to 10µg of DNase I-digested total RNA were denatured for 5 min at 95 °C in RNA loading dye (95% formamide, 0,1% xylene cyanol, 0,1% bromophenol blue, 10mM EDTA), chilled on ice for 2 min, then separated either on 6% denaturing PAA gels (7M urea) or on 1% agarose gels (native conditions). The RNA was transferred to Hybond-XL membrane (GE Healthcare) by electro-blotting at 50V, for 1h using 1X TBE buffer (10X TBE: 890mM Tris base, 890mM Boric acid, 20mM EDTA), then cross-linked to the membrane by UV crosslinking (120kJ). The membranes were pre-incubated for 1h with 15ml of Roti-Hybri-Quick buffer (Roth) at 42 °C and then the radiolabeled probes were added and incubated ON. The membranes were rinsed with 5X SSC (20X SSC: 3 M sodium chloride, 0.3 M sodium citrate, SSC buffer contains in addition 0.1% SDS) to remove the non-hybridized probe, then washed three times at 42 °C with SSC buffer (15 min with 5X SSC, 15 min with 1X SSC and 15 min with 0.1X SSC). RNA signals were visualized on a phosphoimager (Typhoon Trio-Amersham-Bioscience) and band intensities were quantified using MultiGauge software (Fujifilm).

### Slot blot

The slot blots were generated as described in (Hadjeras et al. 2019). 20 μl of cell extract from fractions collected after sucrose gradient fractionation were denatured in the presence of denaturing buffer (2.2 M formaldehyde, 50% formamide, 0.5 mM EDTA, 10 mM MOPS, 4 mM NaCl) and incubated at 65 °C for 5 minutes. Samples were directly placed in a slot on a nylon membrane by vacuum filtration (Amersham Hybond-XL-GE Healthcare) using a transfer collector (PR648-Hoefer ™ Slot Blot). The RNA present in the deposited extracts was irreversibly fixed to the membrane by ultraviolet treatment (120 kJ/cm^2^). The membranes were subsequently hybridized with radiolabeled oligonucleotide probes specific for the rRNAs as described above for Northern blot.

### 5’ RACE

The 5’ ends of rRNAs were mapped using 5’ RACE (Rapid amplification of cDNA ends) analysis following the protocol described (Argaman et al. 2001). First, total RNA or purified rRNAs with 5’ monophosphate ends were ligated to the 3’ hydroxyl group of an RNA oligonucleotide adapter, followed by reverse transcription with a gene-specific primer (RT primer) and subsequent PCR amplification using a 5’-adapter-specific primer and a gene-specific primer. Briefly, the RNA-adapter ligation was performed overnight at 17 °C in the presence of 0.5-1µg total RNA or purified 5S and 5S* rRNA, 21pmol of RNA adapter (RNA A3), 10 units of T4 RNA ligase (ThermoScientific), 1X RNA ligase buffer containing ATP, 15% DMSO and 20 units of RNase Inhibitor in a 20µl final reaction. After addition of 2 pmol of a reverse transcription primer, the reaction was adjusted to a final volume of 150µl by adding milliQ water (RNase-free). Subsequently, the adapter-ligated RNAs were extracted with 1 volume of phenol-chloroform-isoamyl alcohol (P: C: I) in PLG tubes. The aqueous phase was mixed with 3 volumes of a mixture of ethanol and sodium acetate at pH5, ratio 29: 1, to precipitate the RNAs. The adapter-ligated RNAs were dissolved in 30 μl of milliQ water (RNase-free). 0.25-0.5µg of the adapter-ligated RNAs were converted to cDNA using an RT primer specific for each gene encoding the rRNAs and the Superscript III reverse transcriptase as described above (primer extension). After treatment with 1 unit of RNase H (ThermoScientific), 2µl of the cDNA samples were used as template for a PCR reaction using 1 unit of PHUSION® DNA polymerase (Finnzymes), 1X GC buffer, 0.2mM dNTPs, 3% DMSO, and 1µM of the pair of oligonucleotides: the sense primer DNA b6, which anneals to the RNA-adapter sequence, and an antisense primer that anneals within the gene of interest (5S, 16S or 23S). Following visualization on 3% agarose gels, PCR products were excised, purified, and then sequenced after cloning using the Zero Blunt® TOPO® PCR cloning kit (Invitrogen).

### 3’ RACE

0.5 to 1µg total or purified RNA were first dephosphorylated using 1U of thermosensitive Alkaline phosphatase FastAP (ThermoScientific) in the presence of 10X AP buffer and 20U of RNase inhibitor (ThermoScientific) in a final volume reaction of 20µl for 15 minutes at 37 °C. Dephosphorylated RNA was subjected to P:C:I extraction and precipitation with 3 volumes of 30:1 ethanol/sodium acetate solution. The dephosphorylated RNA was ligated to an RNA adapter (RNA E1) ON at 17 °C, P:C:I extracted and precipitated as described above. 0.25 to 0.5µg of the ligated RNA was reverse transcribed in the presence of 5pmol of E4 DNA primer (complementary to the E1 RNA adapter) using 200U of Superscript III (Invitrogen) as described above. After treatment with 1U of RNase H (ThermoScientific), 2µl of the cDNA samples were used as template for a PCR reaction using: 1U of PHUSION® DNA polymerase (Finnzymes), 1X GC buffer, 0.2mM dNTPs, 3% DMSO, and 1µM of a pair of oligonucleotides: the sense primer that anneals within the gene of interest (5S, 16S or 23S), and the antisense primer E4 DNA. Following separation on 3% agarose gels, PCR products were excised, purified, and sequenced after cloning using Zero Blunt® TOPO® PCR cloning kit (Invitrogen).

### cRACE

Purified rRNA fragments extracted from the agarose gel (Fig. S4) were circularized with 20 units T4 RNA ligase (ThermoScientific), 1X RNA ligase buffer containing ATP, 15% DMSO and 20 units of RNase Inhibitor in a 20µl final reaction for 30min at 37 °C. After addition of 2 pmol of a reverse transcription primer, the reaction was adjusted to a final volume of 150µl by adding milliQ water (RNase-free). Subsequently, the circularized RNAs were extracted with 1 volume of phenol-chloroform-isoamyl alcohol (P: C: I) in PLG tubes. The aqueous phase was mixed with 3 volumes of a mixture of ethanol and sodium acetate at pH5, ratio 29: 1, to precipitate the RNAs. 0.25 µg of circularized RNAs were converted to cDNA using an RT primer specific for each gene encoding the rRNAs and the Superscript III reverse transcriptase as described above (primer extension). The reverse transcripts were PCR amplified using PHUSION® DNA polymerase (Finnzymes) and appropriate primers. The products were separated on a 3% agarose gel, purified, and then sequenced after cloning using the Zero Blunt® TOPO® PCR cloning kit (Invitrogen).

### RNA stability experiments

*Escherichia coli* strains were grown on LB at 37 °C to an OD_600_ of 0.4 and then rifampicin was added to a final concentration of 500µg/ml to block new RNA synthesis. Incubation was continued at 37 °C and aliquots were withdrawn at different time points (for example 0, 1, 2, 4, 8, 16, and 32 minutes) after rifampicin addition, mixed with 0.2 volume of stop solution (5% phenol, 95% ethanol v/v), and directly snap frozen in liquid nitrogen. After thawing on ice and pelleting the cells, total RNA was extracted using TRIzol® reagent. RNA levels were measured for the different time points after rifampicin treatment, by primer extension; and the relative half-lives of the 5S* RNAs were calculated. The quantities at different time points are plotted in a semi-logarithmic plot on Microsoft Excel, after normalization by defining the 0 time point (before rifampicin treatment) as having 100% RNA, using exponential fitting. The obtained decay curves appeared linear. The regressions curve function equation was used to determine the relative RNA half-lives.

### Purification of rRNA from polyacrylamide or agarose gels

5 to10 μg of DNaseI-digested total RNA resuspended with the RNA loading buffer (95% formamide, 0.1% xylene cyanol, 0.1% bromophenol blue, 10 mM EDTA), denatured for 5 minutes at 95 ° C, were separated by electrophoresis on 10% polyacrylamide gel in denaturing condition (7 M urea) in 1X TBE buffer (10X TBE: 890mM Tris base, 890mM boric acid, 20mM EDTA) for 5h15 at 300V. To purify 16S and 23S degradation fragments, RNA from sucrose fractions were separated on a 1% agarose gel in 1X TBE buffer for 3h30 at 50V. After staining with SYBR™ Safe stain (Invitrogen) and visualization on ChemiDoc imager (Biorad), the bands corresponding to 5S, 5S* and the different degradation fragments of 16S and 23S were cut and extracted from the gel in 0.3ml RNA elution buffer (0.1 M sodium acetate pH 6.5, 0.1% SDS and 10 mM EDTA pH 8) and incubated with agitation ON at 6-10 °C. After centrifugation at 14000rpm for 15 minutes at 4 °C, RNA was purified using Bio-Spin P30 columns (Bio-Rad). The RNAs were precipitated with 1 volume of absolute ethanol, eluted with 30µl of milliQ water (RNase-free) and then stored a −20 °C.

### Preparation of proteins from sucrose gradient fractions

Sucrose gradient fractions of 0.25 or 0.5 ml were collected, and proteins were precipitated by the addition of trichloroacetic acid (TCA_SIGMA) to a final concentration of 18%. The samples were mixed by inversion and frozen at −20 °C for 30 minutes. The proteins were pelleted by centrifugation at 16000g for 30 minutes at 4 °C and the protein pellets were washed twice with 0.3ml of cold acetone. Acetone was then removed by centrifugation at 16000g for 15 minutes at 4 °C. The proteins were resuspended in 20µl of 20mM Tris-HCl pH 7.5 and then denatured by addition of 2X Laemmli loading buffer (Laemmli 4X (Biorad): 277.8mM Tris-HCl pH6.8, 44.4% glycerol, 4.4% LDS, 0.02% Bromophenol blue) containing 5% β-mercaptoethanol followed by a heating step at 95 ° C for 5 minutes. 10 μl of proteins obtained after TCA precipitation were separated by 4-12% polyacrylamide gradient gel electrophoresis (NuPAGE-Invitrogen) in 1X MES buffer (50 mM MES, 50 mM Tris Base, 0.1% SDS 1 mM EDTA, pH 7.3) for 1h35 at 120V. After electrophoresis, the proteins were transferred to a nitrocellulose membrane (Biorad) using a TransBlot transfer device (Biorad) with the following parameters: mode: heterogeneous molecular weights, 25V, Time, 7 minutes and then subjected to western blotting.

### Western blotting

The blots were treated with anti-L5, anti-S3, anti-L18 polyclonal antibodies provided by Isabelle Iost (INSERM, Bordeaux), hybridized with the second α-sheep-HRP antibody for 1 h at room temperature. Signals were visualized using the ECL kit (Biorad) on a ChemiDoc Imager (Biorad) for chemiluminescence detection.

### Determination of RNase E cleavage site *in vivo* and *in vitro*

To identify the cleavage sites of RNase E, sequences flanking the identified cleavages sites were aligned and analyzed by MEME suite (Version 4. 9. 1) to generate a consensus motif (http://meme-suite.org/) (Bailey et al. 2009) as described (Chao et al. 2017b).

### Ribosomal particles analysis by mass spectrometry

For mass spectrometry analysis, proteins from the different ribosomal particles were prepared in triple biological replicates for each strain. Proteins were extracted from the ribosomal particles after separation on sucrose gradient by adding 18% of cold acetic acid. Protein pellets were resuspended in 25µl of cold 20 mM Tris-HCl, pH 7.5. Protein samples were reduced for 30 min with shaking at 56°C in 2X protein loading buffer (80 mM Tris-HCl pH 6.8, 4% SDS, 20% glycerol, 0.16% BBP, 49.2 mM DTT) and then alkylated in 66 mM iodoacetamide (SIGMA) for 30 min in the dark at room temperature. Equal volumes of the obtained samples were loaded onto 4-12% Bis-Tris Nu-PAGE gel (Thermofisher). For one-shot analysis of the entire mixture, no fractionation was performed, and the electrophoretic migration was stopped as soon as the protein sample migrated for 0.5cm. The gel was briefly stained using then InstantBlue (Expedeon Protein Solutions) according to the manufacturer’s instructions. Each single slice containing the whole sample was excised and subjected to in-gel tryptic digestion using modified porcine trypsin (Promega, France) at 10 ng/μl as previously described (Shevchenko A et al, 2001). The dried peptide extracts obtained were dissolved in 12 μl of 0.05% trifluoroacetic acid in 2% acetonitrile and analyzed by online nanoLC using an Ultimate 3000 RSLCnano LC system (Thermo Scientific Dionex) coupled to an LTQ Orbitrap Velos mass spectrometer (Thermo Scientific, Bremen, Germany) for data-dependent CID fragmentation experiments. 5μl of each peptide extracts were loaded in two or three injection replicates onto 300μm ID x 5mm PepMap C18 precolumn (ThermoFisher, Dionex) at 20 μl/min in 2% acetonitrile, 0.05% trifluoroacetic acid. After 5 minutes of desalting, peptides were online separated on a 75 μm ID x 50 cm C18 column (in-house packed with Reprosil C18-AQ Pur 3 μm resin, Dr. Maisch; Proxeon Biosystems, Odense, Denmark), equilibrated in 95% of buffer A (0.2% formic acid), with a gradient of 5 to 25% of buffer B (80% acetonitrile, 0.2% formic acid) for 80min then 25% to 50% for 30 min at a flow rate of 300 nL/min. The LTQ Orbitrap Velos was operated in data-dependent acquisition mode with the XCalibur software (version 2.0 SR2, Thermo Fisher Scientific). The survey scan MS was performed in the Orbitrap on the 350–1,800 m/z mass range with the resolution set to a value of 60,000. The 20 most intense ions per survey scan were selected with an isolation width of 2 m/z for subsequent data-dependent CID fragmentation and the resulting fragments were analyzed in the linear trap (LTQ). The normalized collision energy was set to 30%. To prevent repetitive selection of the same peptide, the dynamic exclusion duration was set to 60 s with a 10 ppm tolerance around the selected precursor and its isotopes. Monoisotopic precursor selection was turned on. For internal calibration the ion at 445.120025 m/z was used as lock mass.

### Database search and label-free quantitative analysis

All raw MS files were processed with MaxQuant (v 1.6.1.0) for database search with the Andromeda search engine and for quantitative analysis. Data were searched against the UniProtKB/Swiss-Prot protein database released 2018_04 with *Escherichia coli* (K12 strain) (5,979 sequences) supplemented with a list of frequently observed contaminant sequences provided in MaxQuant. Enzyme specificity was set to trypsin/P, and a maximum of two missed cleavages was allowed. Carbamidomethylation of cysteines was set as a fixed modification, whereas methionine oxidation was set as variable modification. The precursor mass tolerance was set to 20 ppm for the first search and 10 ppm for the main Andromeda database search, and the mass tolerance in MS/MS mode was set to 0.8Da. The required minimum peptide length was seven amino acids, and the minimum number of unique peptides was set to one. Andromeda results were validated by the target-decoy approach using a reverse database and the false discovery rates at the peptide-spectrum matches (PSM) and protein level were set to 1%. For label-free relative quantification of the samples, the match between runs option of MaxQuant was enabled with a time window of 2 min, to allow cross-assignment of MS features detected in the different runs after alignment of the runs with a time window of 20 min. Protein quantification was based on razor peptides. The minimum ratio count was set to 1 for label-free quantification calculation, and computation of the intensity based absolute quantification (iBAQ) metric was also enabled.

To perform relative quantification between all identified proteins we used the normalized “LFQ intensity” metric from the MaxQuant “proteinGroups.txt” output. Protein groups with negative identification scores were filtered, as well as proteins identified as contaminants. After log2-transformation of LFQ intensities, log transformed protein intensities corresponding to different technical LC-MS replicate runs were averaged and missing values were replaced to a mean LFQ intensity value was computed from technical LC-MS replicate runs by a noise value randomly drawn using the Perseus software (version 1.5.3.0). For each pairwise comparison of protein content of the subparticles 20S and 40S with their parental 30S and 50S from *rne*ΔMTS and with the 30S and 50S from *rne*+, an unpaired two-tailed Student’s t-test was performed based on the protein intensities. Proteins were considered significantly enriched when their absolute log2-transformed fold change was higher than 1 and their p-value lower than 0.05. To eliminate false-positive hits from quantitation of low intensity signals, two additional criteria were applied: only the proteins identified with a total number of averaged peptide spectrum match (PSM) counts>4 and quantified in a minimum of two biological replicates, before missing value replacement, for at least one of the two compared conditions were selected. Volcano plots were drawn to visualize significant protein abundance variations between the compared ribosomal particles. They represent −log10 (p-value) according to the log2 ratio. The complete list of the identified and quantified proteins and analyzed according to this statistical procedure is described in Table S3.

### *In vitro* cleavage assay

Expression and purification of RNase E(1-598) with a C-terminal HISx6 tag in BL21(DE3) was as described (Khemici et al. 2008). Briefly, cells were lysed and debris were removed by centrifugation at 10,000 g, 4° C for 1 h. The cleared lysate was applied to an NTA-Ni column and eluted with an imidazole gradient. Peak fractions were dialyzed overnight in storage buffer (10 mM Tris HCl, pH 7.5 - 500 mM NaCl - 50% glycerol - 0.2% Genapol X-080 - 10 mM MgSO_4_ - 1 mM EDTA - 1 mM TCEP - 1x Protease Inhibitor), flash frozen in liquid N_2_. and stored at −80° C. Concentration was determined by UV absorbance at 280 nm using a molar extinction coefficient calculated from the amino acid composition of the protein. Ribosomes were purchased from NEB (P0763S). Ribosomal RNA was prepared by extraction of ribosomes using a Direct-zol™ RNA purification kit.

The RNase E cleavage assay was performed in a total reaction volume of 25 µl in either high ionic strength buffer (70 mM Tris, pH7.5, 100 mM KCl, 25 mM MgCl_2_, 10 mM DTT, 2U RNase inhibitor) or low ionic strength buffer (70 mM Tris, pH7.5, 10 mM DTT, 2U RNase inhibitor). Ribosomes (0.22 µM), or a comparable amount of rRNA (16 µg) were incubated with 0.3 µM RNase E(1-598)-Hisx6 at 37° C for 0, 30 or 60 min at 37 °C. Reactions were quenched with 75 µl of cold 10 mM EDTA and held on ice. RNA was extracted using Direct-zol™ RNA purification kit, eluted in 60 µl water (RNase-free), lyophilized, and suspended in 10 µl of RNA loading dye (95% formamide, 0,1% xylene cyanol, 0,1% bromophenol blue, 10mM EDTA). The sample was incubated for 3 min at 95 °C and then separated on a 1% agarose gel in 1x TBE for 195 min at 50V. The gel was stained with SYBR™ Green stain (Invitrogen).

## Acknowledgements

This work was supported by grants from the French National Research Agency (ANR-13-BSV6- 0005; ANR-16-CE12-0014-02). LH was awarded a predoctoral fellowship from the French Ministry of Education. The work was also supported in part by the French Ministry of Research with the Investissement d’Avenir Infrastructures Nationales en Biologie et Santé program (ProFI, Proteomics French Infrastructure project, ANR-10-INBS-08). We thank Isabelle Iost for antibodies against ribosomal proteins.

## Author contributions

LH, MB and AJC initiated the project. LH, MB, LP, CF, OBS, LH, LG, MCB and AJC designed experiments and analyzed data. LH, MB, IC, LP, QMO, CF, LH and LG performed experimental work. LH and AJC wrote the manuscript with feedback from the other authors.

## Declaration of interest

The authors declare that they have no conflict of interest.

---

**Table 1.** Strains and plasmids.

| Strain | Genotype | Reference |
| --- | --- | --- |
| NCM3416 | *E. coli* K12, F<sup>-</sup>, λ<sup>-</sup>, *rph*<sup>+</sup>, *zib-207::Tn10* | (Soupene et al. 2003) |
| MG1655 | F<sup>-</sup>, λ<sup>-</sup>, *rph*<sup>-</sup> | (Jensen 1993) |
| MBS106 | NCM3416, *rne-frt* | (Hadjeras et al. 2019) |
| MBS157 | NCM3416, *rneΔMTS-frt* | (Hadjeras et al. 2019) |
| BL21(DE3) | F<sup>-</sup>, *ompT*, *hsdS<sub>B</sub>* (*r<sub>B</sub><sup>-</sup>*, *m<sub>B</sub><sup>-</sup>*), *dcm*, *gal*, λ(DE3) | (Studier and Moffatt 1986) |
| Kti658 | MG1655, *rneΔMTS-frt* | (Hadjeras et al. 2019) |
| Kti665 | NCM3416 *rne*<sup>+</sup> Δ*pcnB* | This work. |
| Kti669 | NCM3416 *rneΔMTS* Δ*pcnB* | This work. |
| LHS-457 | NCM3416 *rne*<sup>+</sup> Δ*pcnB* Δ*pnp* | This work. |
| LHS-459 | NCM3416 *rneΔMTS* Δ*pcnB* Δ*pnp* | This work. |
| SQM-5 | NCM3416 *rne*<sup>+</sup> Δ*pnp* | This work. |
| SQM-6 | NCM3416 *rneΔMTS* Δ*pnp* | This work. |
| SLP-35 | NCM3416 *rne*<sup>+</sup> Δ*rnr* | This work. |
| SLP-36 | NCM3416 (Huang et al. 2020) *rneΔMTS* Δ*rnr* | This work. |
| CA244 | *rnr*<sup>-</sup> *pnp-200* *Cm<sup>r</sup>* | (Cheng and Deutscher 2003) |
| SLP-79 | NCM3416 *rne*<sup>+</sup> Δ*rnr* *pnp200-kn* | This work. |
| SLP-80 | NCM3416 *rneΔMTS* Δ*rnr* *pnp200-kn* | This work. |
| Kti162 | NCM3416, *rne-mcherry* | (Strahl et al. 2015) |
| Kti513 | NCM3416, *rneΔMTS-mcherry* | (Strahl et al. 2015) |
| Plasmid | Features | Reference |
| pLP56-2 | pET-*rne*(1-598)-his6 | (Hadjeras et al. 2019) |

**Table 2.**
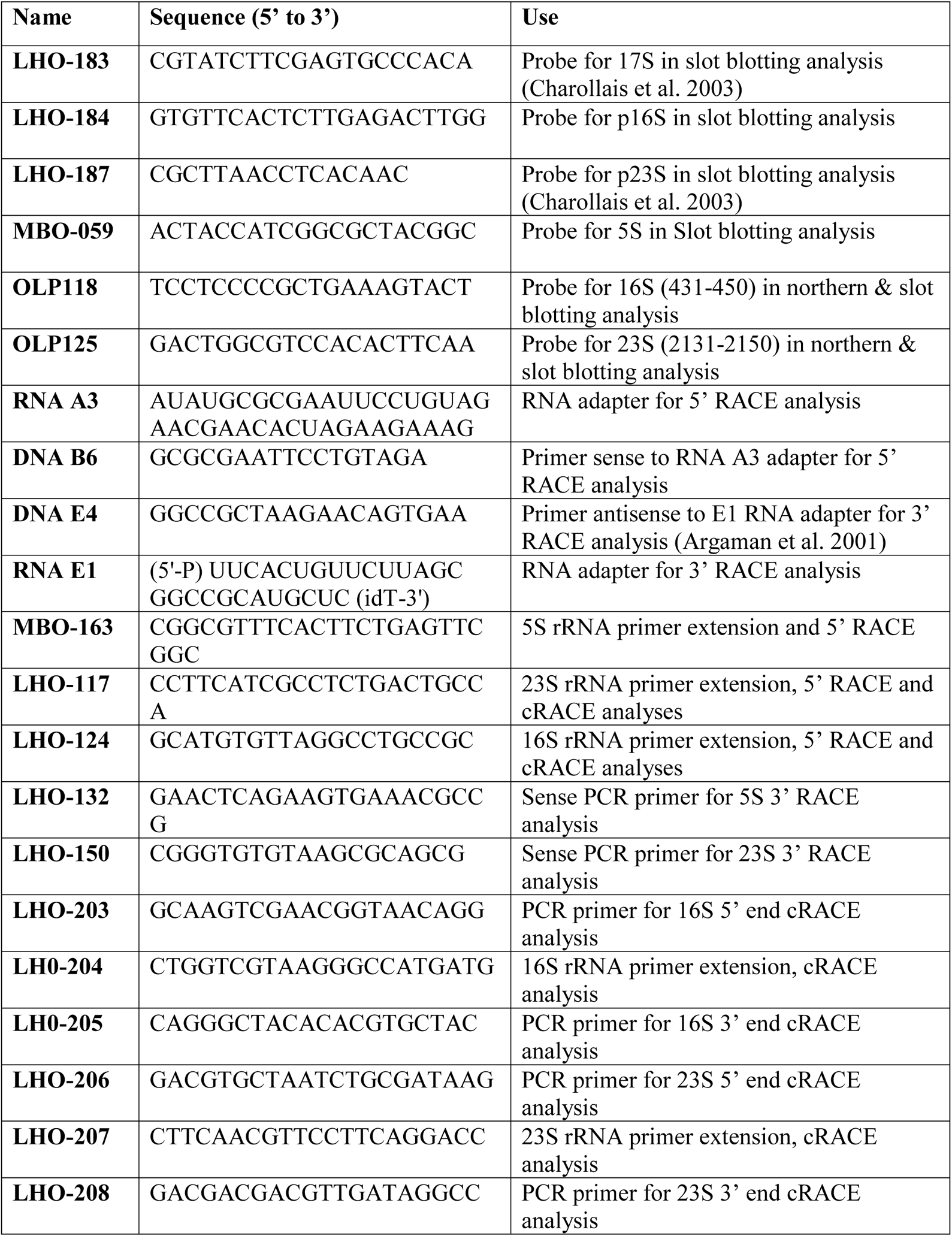
Primers.

**Table S1. cRACE data.** (Excel file)

**Table S2. Mass spectrometry data.** (Excel file)

## Figures with legends

**Fig. S1.**
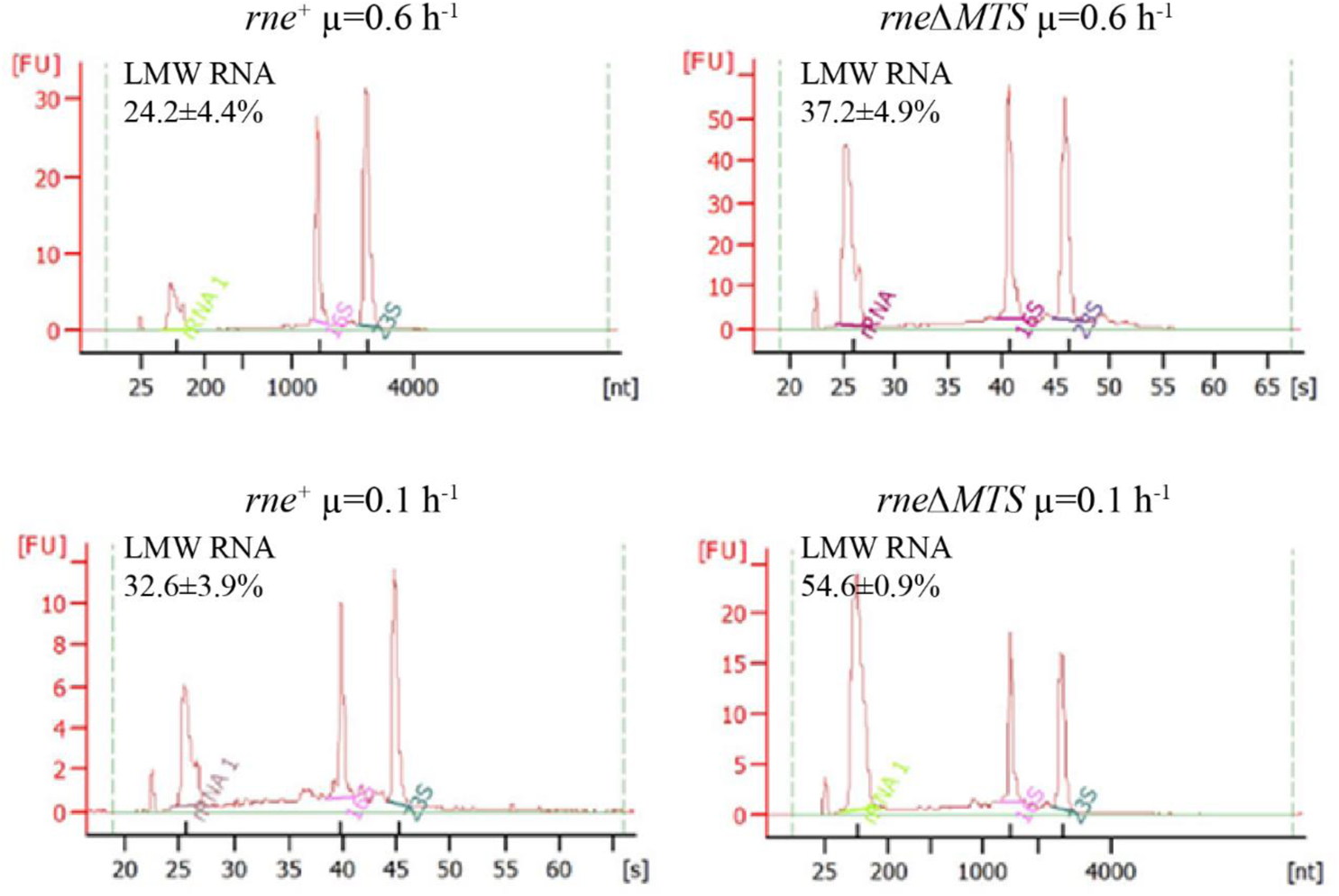
Bioanalyzer analyses of total RNA preparations. Electrophoretograms of total RNA isolated from cultures grown in minimal glucose medium at fast (µ=0.6 h^-1^) and slow (µ=0.1 h^-1^) growth rates. RNA levels were measured by fluorescence (FU) and elution was expressed either as seconds (s) or size (nt). The level of RNA in the peak centered at 100 nt was quantified as the percentage of total RNA. Under both fast and slow growth conditions, there was an approximately 60% increase in the level of Low Molecular Weight (LMW) RNA in the *rne*ΔMTS strain suggesting an accumulation of RNA degradation products.

**Fig. S2.**
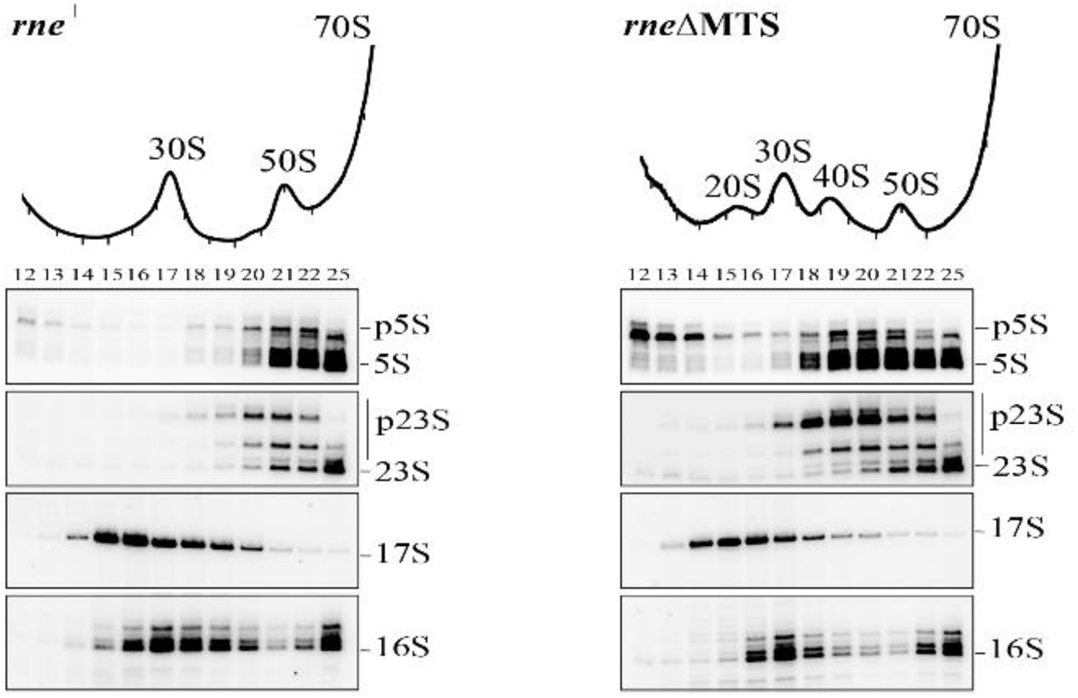
Ribosomal RNA 5’ end mapping by primer extension. Equal volume of clarified cell lysates from *rne*^+^ (left) and *rne*ΔMTS (right) strains were fractionated by velocity sedimentation under condition that optimized separation in the range of 5S to 70S. RNA from the sucrose gradient fractions was analyzed by primer extensions using [^32^P] end-labelled oligonucleotides specific to the 5’ ends of 5S, 23S, 17S and 16S rRNA. After extension by reverse transcriptase, the products were separated by denaturing gel electrophoresis. The 5’ end of mature rRNA and that of the prominent precursors are indicated to the right of each panel. Band located between the p5S and 5S ends correspond to the 5S+1 and 5S+2 species.

**Fig. S3.**
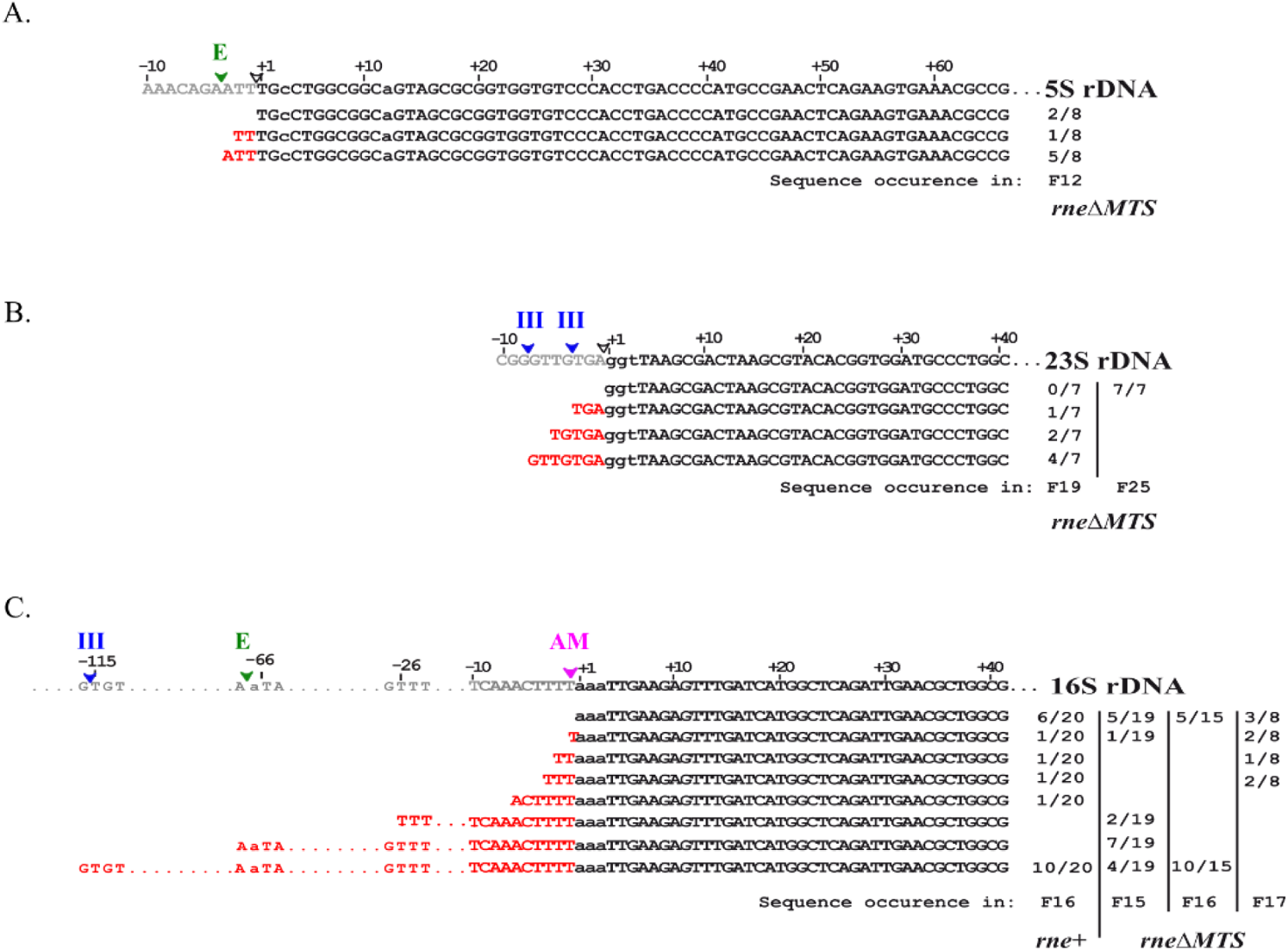
Mapping of rRNA 5’ ends by RACE. RNA from sucrose gradient fractions (Fig. 2A) was analyzed by linker ligation to the RNA 5’ end, PCR amplification and DNA sequencing (5’ RACE). 5’ ends were aligned with the sequence of 16S or 23S rRNA from the *E. coli rrfB* operon. 5’ extensions are indicated in red. Ribosomal RNA processing sites are indicated by arrows: RNase III, blue; RNase E, green; RNase AM, pink. The number of times each sequence was detected is indicated on the right. **A.** 5S rRNA, **B.** 23S rRNA and **C.** 16S rRNA.

**Fig. S4.**
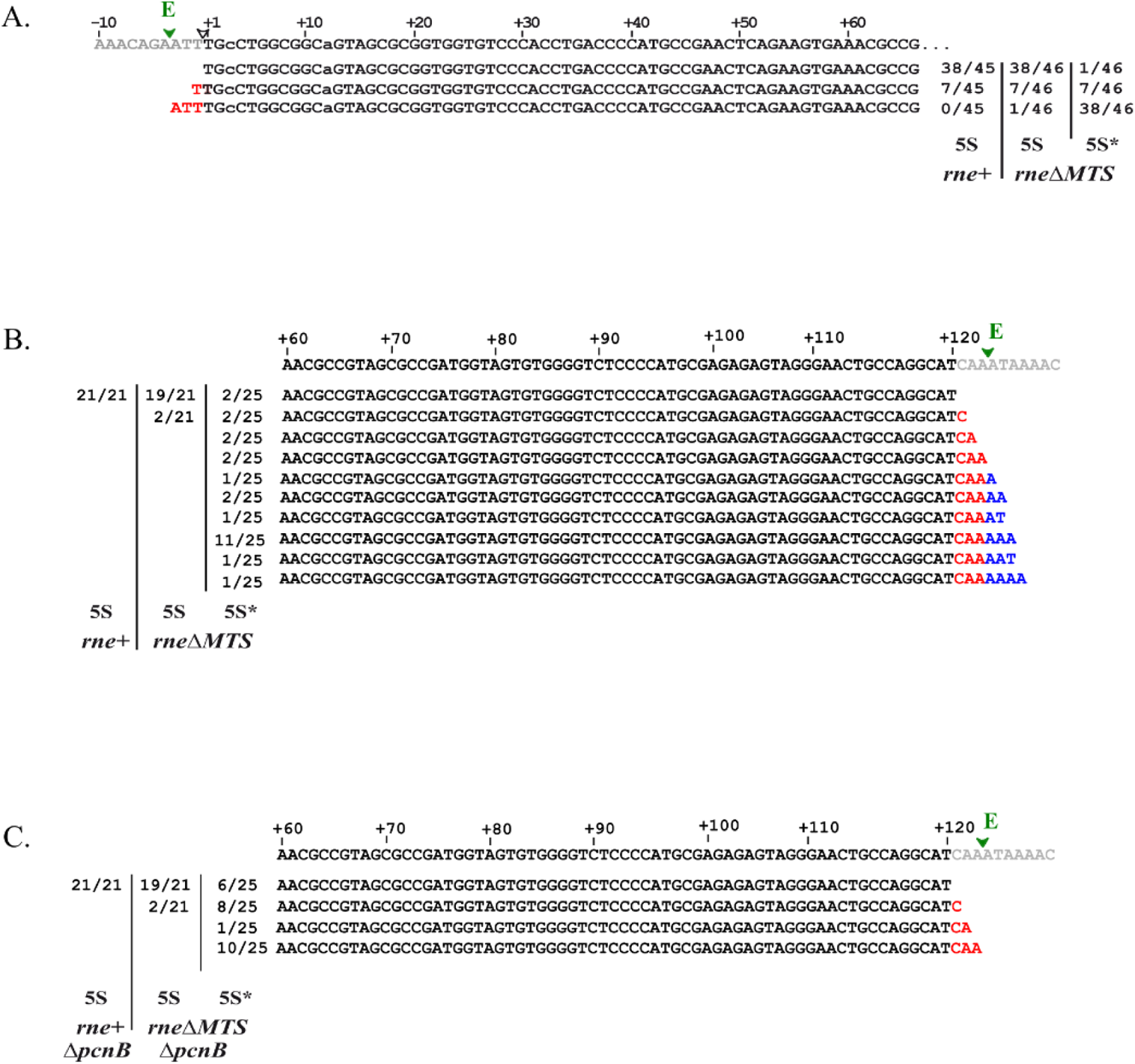
Mapping of 5S and 5S* 3’ ends by RACE. 5S and 5S* rRNA species were extracted from the gel shown in Fig. 3A and subjected to 5’ and 3’RACE analysis. Reference sequences are from the *E. coli rrfB* operon. 5’ and 3’ extensions are indicated in red; untemplated extension in blue. RNase E processing sites are indicated by the green arrow. The occurrence for each sequence is summarized on the right (3’ RACE) or left side (5’ RACE). **A.** 5’ end analysis of 5S* rRNA. 5’ RACE on 5S and 5S* rRNAs from *rne*ΔMTS strain. For comparison, 5S rRNA from *rne*^+^ strain was processed in parallel. **B.** 3’ end analysis of 5S* rRNA. 3’ RACE on 5S and 5S* rRNAs from *rne*ΔMTS strain. For comparison, 5S rRNA from *rne*^+^ strain was processed in parallel. **C.** As in **B.** except in the *ΔpcnB* strain background.

**Fig. S5.**
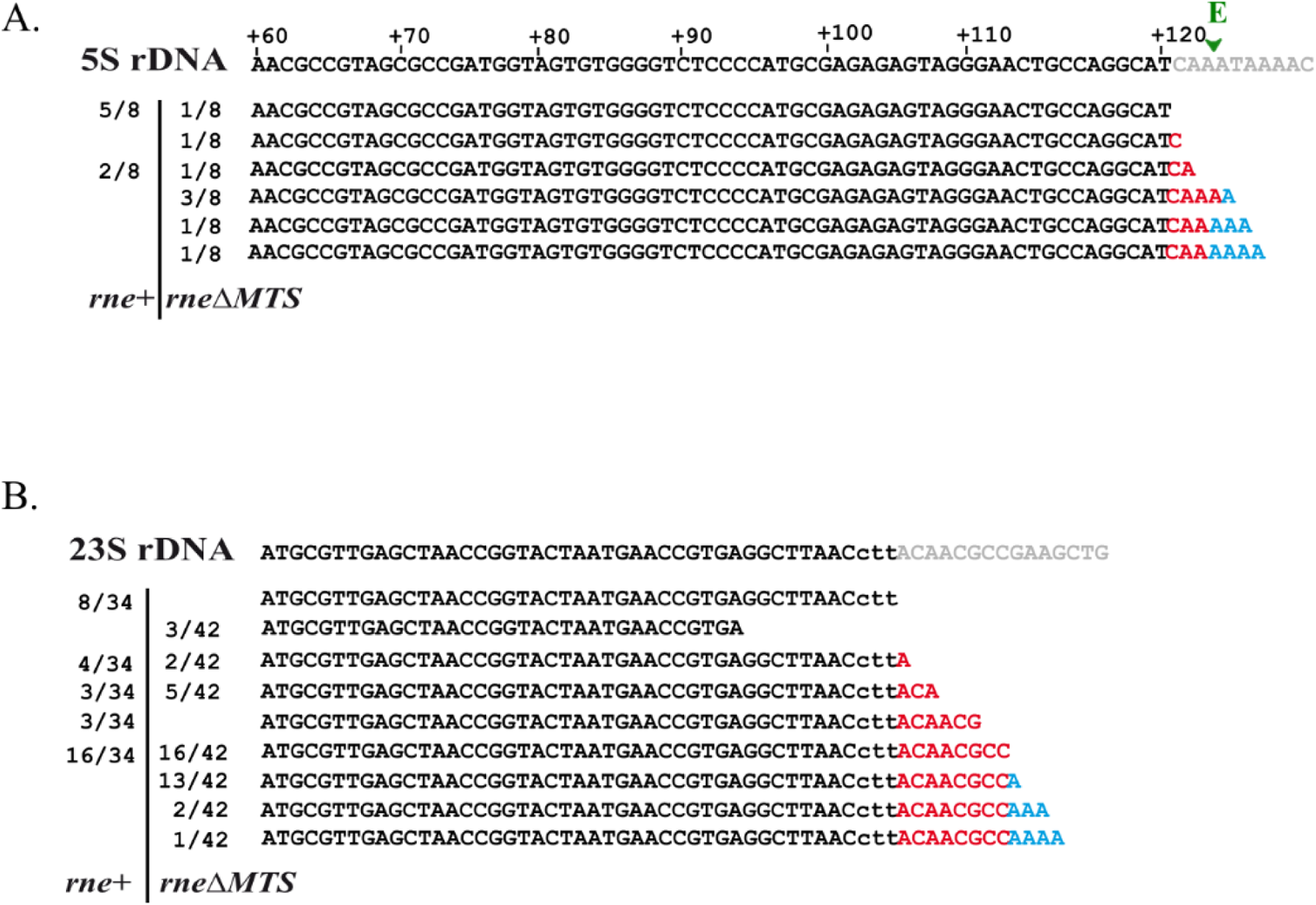
Mapping of rRNA 3’ ends by RACE. RNA from sucrose gradient fractions (Fig. 2A) was analyzed by linker ligation to the RNA 3’ end, PCR amplification and DNA sequencing (3’ RACE). 3’ ends were aligned with the sequence of 5S or 23S rRNA from the *E. coli rrfB* operon. 3’ extensions are indicated in red; untemplated A additions in blue. 5S rRNA (**A.**) and 23S rRNA (**B.**) from the 50S subunit (*rne^+^*) and 40S particle (*rneΔMTS*).

**Fig. S6.**
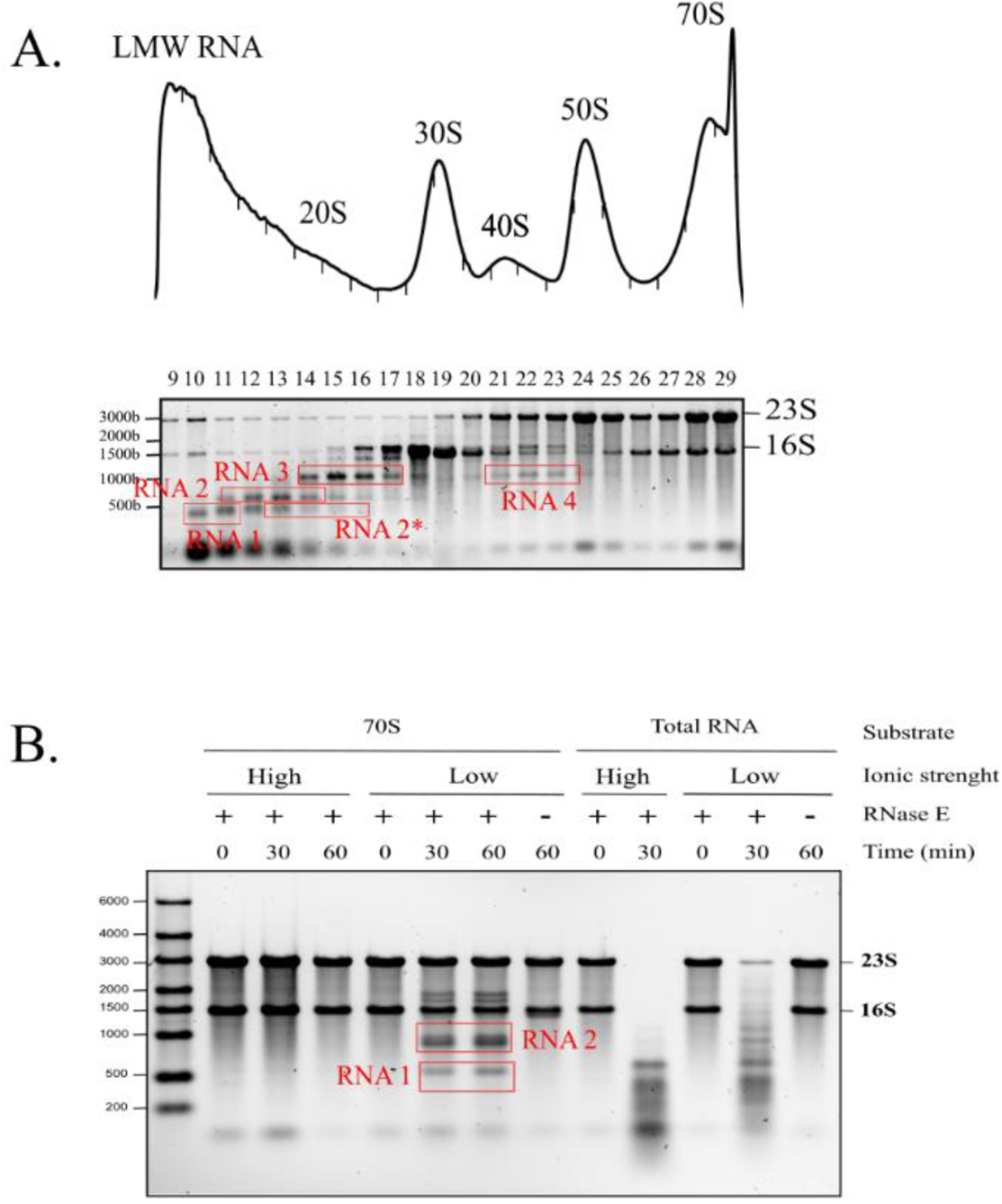
Analysis of rRNA fragments by cRACE. Markup of gels shown in Fig. 4A showing bands that were excised for RNA extraction and cRACE analysis. After RNA circularization, cDNA copies corresponding to the junction of the 5’ and 3’ ends were gel purified and cloned into a plasmid vector. The 5’-3’ ends were then identified by sequencing the cloned cDNA fragments. **A.** *In vivo* fragments. **B.** *In vitro* fragments.

## Notes

### Competing Interest Statement

The authors have declared no competing interest.

### Summary of Updates

The pre-print has been revised to improve the text. Tables and figures are the same as in the first version.

